# Biochemical fractionation of human α-Synuclein in a *Drosophila* model of synucleinopathies

**DOI:** 10.1101/2024.02.05.579034

**Authors:** Khondamir Imomnazarov, Joshua Lopez-Scarim, Ila Bagheri, Valerie Joers, Malú Gámez Tansey, Alfonso Martín-Peña

## Abstract

Synucleinopathies are a group of central nervous system pathologies that are characterized by neuronal accumulation of misfolded and aggregated α-synuclein in proteinaceous depositions known as Lewy Bodies (LBs). The transition of α-synuclein from its physiological to pathological form has been associated with several post-translational modifications such as phosphorylation and an increasing degree of insolubility, which also correlate with disease progression in post-mortem specimens from human patients. Neuronal expression of α-synuclein in model organisms, including *Drosophila melanogaster,* has been a typical approach employed to study its physiological effects. Biochemical analysis of α-synuclein solubility via high-speed ultracentrifugation with buffers of increasing detergent strength offers a potent method for identification of α-synuclein biochemical properties and the associated pathology stage. Unfortunately, the development of a robust and reproducible method for evaluation of human α-synuclein solubility isolated from *Drosophila* tissues has remained elusive. Here, we tested different detergents for their ability to solubilize human α-synuclein carrying the pathological mutation A53T from brains of aged flies. We also assessed the effect of sonication on solubility of human α-synuclein and optimized a protocol to discriminate relative amounts of soluble/insoluble human α-synuclein from dopaminergic neurons of the *Drosophila* brain. Our data established that, using a 5% SDS buffer, the 3-step protocol distinguishes between cytosolic soluble proteins in fraction 1, detergent-soluble proteins in fraction 2 and insoluble proteins in fraction 3. This protocol shows that sonication breaks down α-synuclein insoluble complexes from the fly brain, making them soluble in the SDS buffer and enriching fraction 2 of the protocol.

## INTRODUCTION

Misfolding and aggregation of the product encoded in the human α-synuclein gene (*hSNCA*) characterizes a group of nervous system disorders known as synucleinopathies (1, 2). These disorders include Parkinson’s disease (PD), the second most common neurodegenerative disease, PD dementia (PDD), multiple system atrophy (MSA), Lewy body dementia (LBD) and Dementia with Lewy Body (DLB), among others (1, 2), many of which are classified under the umbrella term of parkinsonism due to their effects on motor control (slowed movement, rigidity, and/or tremor). A common pathological hallmark of synucleopathies is the deposition of α-synuclein in intracellular neuronal inclusions termed Lewy Bodies (LB).

Pathological α-synuclein is the major protein component of LBs, where it is mainly found in its insoluble fibrillar form. Conversely, in neurons, physiological α-synuclein is located in the cytoplasm and/or is associated with membranous structures in synaptic terminals in a soluble monomeric form (3). Although monomeric α-synuclein can aggregate spontaneously to form amyloid fibrils, which are the core components of LBs, α-synuclein presents a stable conformation upon binding to lipid membranes (4). The transition of α-synuclein from a benign soluble form into the amyloidogenic fibrillar conformation that constitutes the principal scaffold component of LBs is thought to be caused by extensive post-translational modifications (5–10), rather than a mere increase in its concentration (4, 11). Though protein abundance positively correlates with a higher propensity for aggregation (12–14). Furthermore, the presence of LB and, consequently, histopathologic deposition of α-synuclein occurs in 80-90% of PD cases.

In general, increasing insolubility of α-synuclein and a decrease in soluble α-synuclein in the frontal cortex of postmortem PD brains were reported to positively correlate with both disease duration and disease stage (15). Postmortem samples of the basal ganglia and the limbic cortex from patients with idiopathic PD also show a drastic increase in insoluble α-synuclein content compared to age-matched healthy controls (16).

Given the major role that α-synuclein plays in the pathophysiology of these neurodegenerative diseases, the ability to express human α-synuclein in the fruit fly, *Drosophila melanogaster*, represents an invaluable system to investigate its effects on brain physiology, behavior, and disease progression, in particular PD. The genetic amenability and tractability of the fly have provided considerable insights into PD pathogenesis (17–34) demonstrating, for instance, the selective vulnerability of dopaminergic neurons to α-synuclein toxicity (22), the protective role of glucocerebrosidase against α-synuclein aggregation (19, 27), the importance of VPS35 for lysosomal α-synuclein degradation(26), or the protective activity of DJ-1 against oxidative stress (25). The original work expressing *hSNCA* in the fly brain provided histological evidence of α-synuclein aggregation, revealing punctate immunostaining characteristic of proteinaceous aggregates, and electron micrographs illustrating α-synuclein inclusions with radiating protein filaments in neurons (22). However, a robust biochemical characterization of these α-synuclein inclusions in *Drosophila* models has been lacking; therefore, the extent to which these α-synuclein aggregates resemble those found in LBs from human brain specimens, remains unclear. Given that the progression in the pathophysiology of these diseases strongly depends on the transition from monomeric soluble α-synuclein to increasingly insoluble species (15), it becomes critical to develop a robust protocol to identify the specific α-synuclein species and their solubility status at each stage of the model organism’s lifespan. Developing the tools and techniques for this identification is instrumental to unravel the molecular mechanisms that underlie the transition from physiological soluble monomers to pathological insoluble aggregates or to detect therapeutic effects on α-synuclein assemblies for future physiological, behavioral, or treatment studies in *Drosophila* models of synucleopathies. Here we implemented a multi-step fractionation protocol in which the second fraction employs a buffer containing 5% SDS that solubilizes membrane-associated α-synuclein but not insoluble α-synuclein. The combination of this buffer with a 3-step serial fractionation procedure exhibits such resolution that we could detect changes in α-synuclein segregation upon sonication of the samples prior to the fractionation protocol, also demonstrating that the aggregates formed in DA neurons of *Drosophila* can be disaggregated upon this mechanical application.

## RESULTS

### Differential solubility of α-synuclein in selective detergents

Disease progression largely correlates with species-specific (from monomers to oligomers and fibrils) accumulation of α-synuclein and its deposition into insoluble neuronal inclusions (LBs) (15). Chemical fractionation of human specimens or tissues from vertebrate models has provided benchmark methods for quantification of α-synuclein solubility and identification of pathology stage. To translate this methodology to *Drosophila* models of synucleinopathies, we first tested different buffers containing selective detergents for their ability to fractionate human α-synuclein (hSNCA) expressed in dopaminergic (DA) neurons of the fly brain. Flies were aged up to 20 days post-eclosion at 25C to allow time for α-synuclein aggregation; by this time point, flies have developed significant behavioral and cellular phenotypes (22). At this age, we snap-froze flies in liquid nitrogen and collected heads from controls (*TH-Gal4/UAS-LacZ*) and α-synuclein expressing flies (*TH-Gal4/UAS-hSNCA^A53T^*). We then homogenized and sonicated the samples prior to the procedure for sequential protein extractions. The multi-step fractionation protocol resulted in three fractions where different detergent solvents were tested in the second fraction: the first fraction (fraction 1: TBS-soluble) extracted TBS-soluble proteins; the second fraction (fraction 2: detergent-soluble) extracted either TBS-soluble (TBS-wash), SDS-soluble, or RIPA-soluble proteins; and, finally, we sonicated and resuspended the TBS-insoluble, SDS-insoluble or RIPA-insoluble pellet in urea and SDS (fraction 3: insoluble) (Fig. 2A). We detected similar amounts of α-synuclein in the TBS-soluble fractions (fraction 1) from these three protocols (Fig. 2B-C). However, in the second fraction, we only detected α-synuclein when solubilizing protein extracts in the SDS (SDS-soluble) or RIPA (RIPA-soluble) buffers, with a significantly higher yield in RIPA-soluble fraction (Fig. 2B-C). The second TBS fraction (fraction 2: TBS-wash) was unable to extract any α-synuclein, indicating that the TBS-soluble α-synuclein was fully extracted in the first fraction. The insoluble fractions (fraction 3) were resuspended in urea/SDS and we observed low but detectable amounts of α-synuclein in the protocols that used the SDS buffer (Table 1) for the second fraction (Fig. 2B-C). However, we were not able to consistently detect α*-*synuclein in the insoluble fraction that followed RIPA extraction (fraction 3: RIPA-insoluble; Fig. 2B-C), being detectable in 2 of 5 experiments. Together, these data indicate that expression of *hSNCA* in DA neurons of the fruit fly, *Drosophila melanogaster*, led to accumulation of TBS-insoluble α-synuclein, that was most abundant in the detergent-containing fraction (SDS- and RIPA-soluble fraction 2) of the protocol (Table S2). In addition, the brain-accumulated α-synuclein exhibited sufficient high solubility in RIPA buffer that no additional protein was detectable in the insoluble (urea) fraction.

**Figure 1.**
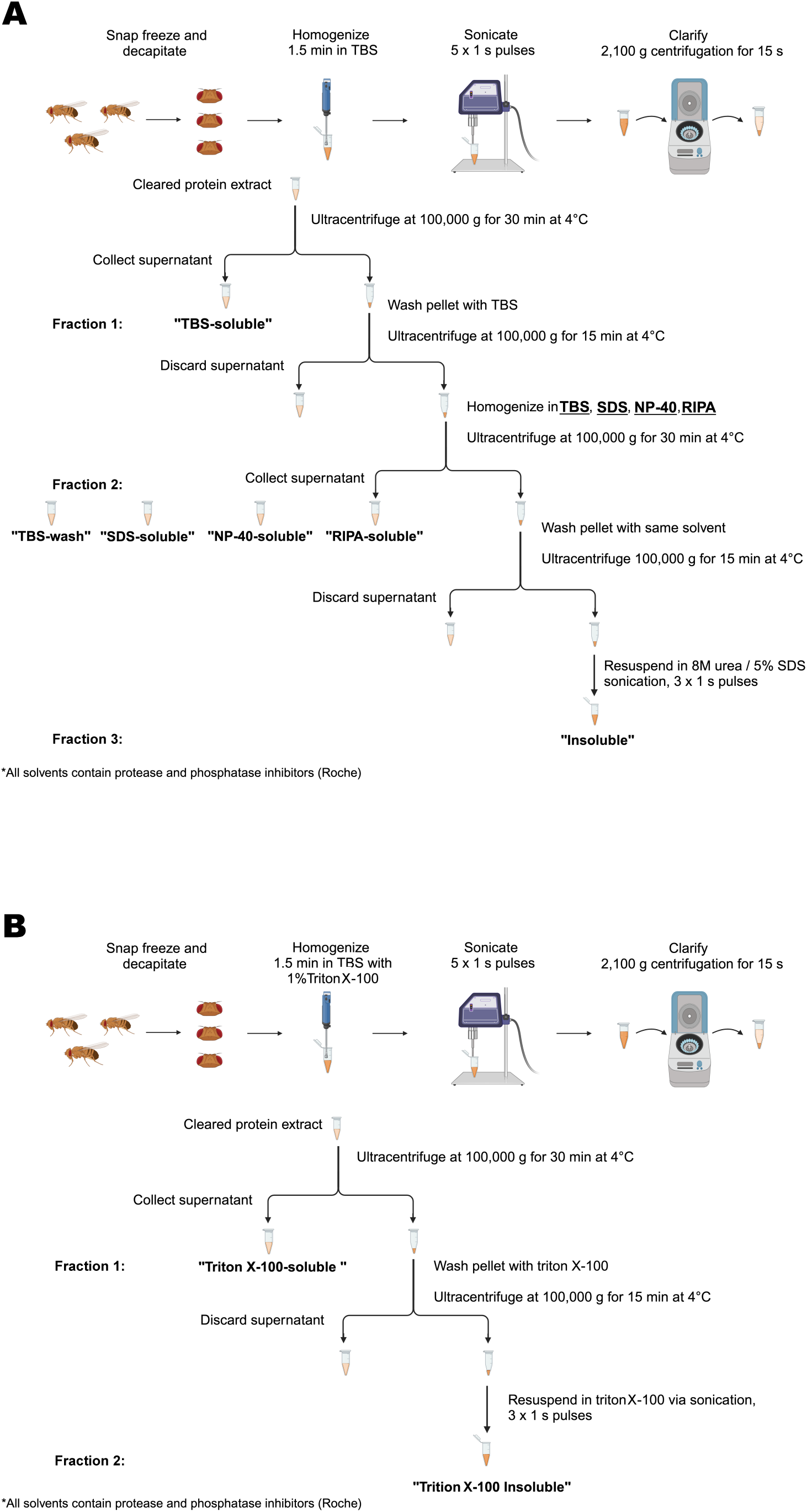
Schematic representation of the procedure for serial protein fractionation from *Drosophila* heads. **(A)** A 3-step sequential ultracentrifugation protocol for extracting insoluble α-synuclein from fly heads using TBS, SDS, RIPA or NP-40 buffers. (B) A 2-step sequential ultracentrifugation protocol for extraction of α-synuclein from fly heads using Triton X-100 as a solvent in the fractionation buffer.

**Figure 2.**
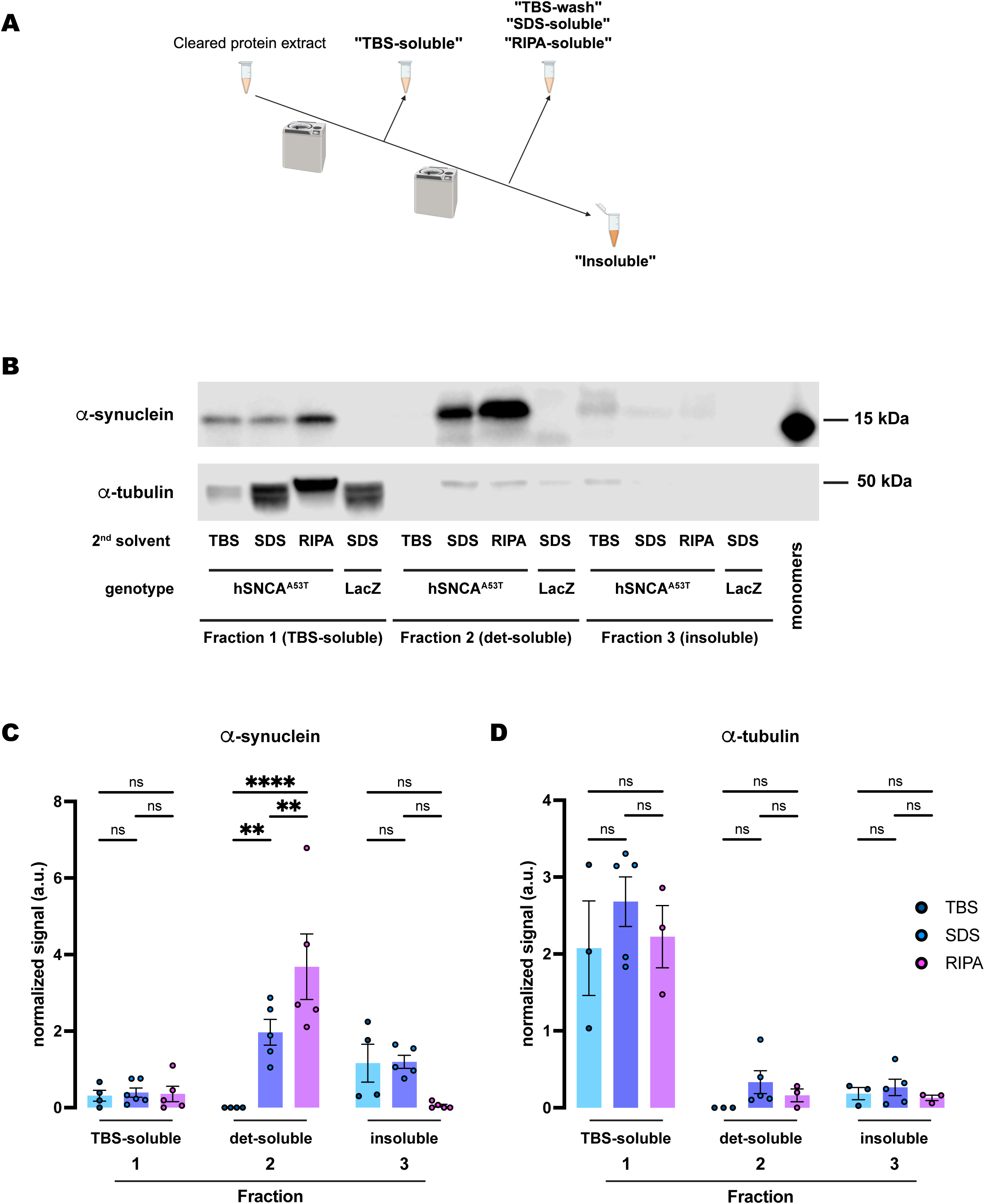
Human A53T α-synuclein extracted from *Drosophila* dopaminergic neurons exhibits higher solubility in RIPA than in the SDS buffer. **(A)** Schematic representation of the sequential fractionation protocol and the extraction buffers employed in this experiment. **(B)** Representative western blot of head lysates from flies expressing *hSNCA^A53T^*in dopaminergic neurons. Fly heads are fractionated using a 3-step protocol in which the second fraction uses a variable detergent solvent, TBS, SDS or RIPA buffer. The first fraction (TBS-soluble) is loaded in lanes 1-4, second fraction (TBS-wash, SDS-soluble or RIPA-soluble) is loaded in lanes 5-8, and the third fraction (insoluble) is loaded in lanes 9-12, while 2ng of purified recombinant human α-synuclein monomers (monomer) are loaded in lane 13 as positive control. Protein lysates are extracted from flies expressing *hSNCA^A53T^* in dopaminergic neurons (*w; +/+; TH-Gal4/UAS-hSNCA^A53T^*, lanes 1-3, 5-7, 9-11) and control flies (*w; +/+; TH-Gal4/UAS-LacZ*) as negative controls not expressing hSNCA (lanes 4, 8, 12). The fractions are probed for α-synuclein (4B12, top panel) and α-tubulin (T6074, bottom panel). **(C)** Quantification of α-synuclein content across independent experiments shows no significant differences between the different solvents in the TBS-soluble fraction (Tukey’s multiple comparisons; TBS vs SDS, p=0.9864; SDS vs RIPA, p=0.9972; TBS vs RIPA, p=0.9959) or the insoluble fraction (Tukey’s multiple comparisons; TBS vs SDS, p=0.7746; SDS vs RIPA, p=0.3137; TBS vs RIPA, p=0.1097) but significantly more α-synuclein is detected in the SDS- and RIPA-soluble fractions than in the TBS-wash (Tukey’s multiple comparisons; TBS vs SDS, p=0.0022; SDS vs RIPA, p=0.0048; TBS vs RIPA, p<0.0001). **(D)** Quantification of α-tubulin content shows no significant differences between the solvents (Tukey’s multiple comparisons; Fraction 1: TBS vs SDS, p=0.2462; SDS vs RIPA, p=0.4421; TBS vs RIPA, p=0.9298; Fraction 2: TBS vs SDS, p=0.6415; SDS vs RIPA, p=0.8869; TBS vs RIPA, p=0.9189; Fraction 3: TBS vs SDS, p=0.9736; SDS vs RIPA, p=0.9267; TBS vs RIPA, p=0.9899). Samples size was n=4 (α-synuclein) and n=3 (α-tubulin) for TBS samples, n=5 (α-synuclein and α-tubulin) for SDS samples, and n=5 (α-synuclein) and n=3 (α-tubulin) for RIPA samples. Error bars indicate SEM. Tukey’s comparison test: n.s. not significant, * p<0.05, ** p<0.005, *** p<0.001.

**Table 1:** Chemical composition of the solutions for biochemical fractionation. Solutions and buffers employed in this study are listed and their chemical composition specified. Note that the SDS buffer only contains SDS, while the RIPA buffer contains SDS at a much lower concentration in addition to NP-40 and sodium deoxycholate.

| Name | Composition | Manufacturer (catalog number) |
| --- | --- | --- |
| TBS | 150 mM NaCl, 20mM Tris base, pH=7.6 | Made in-house |
| SDS | 5% w/v sodium dodecyl sulfate (SDS), 150 mM NaCl, 20mM Tris base, pH=7.6 | Made in-house |
| RIPA | 1% NP-40, 1% sodium deoxycholate, 0.1% SDS, 150 mM NaCl, 25 mM Tris HCl, pH=7.6 | Thermo Fisher Scientific (89900) |
| NP-40 | 1% NP-40 (74385, Sigma-Aldrich), 150 mM NaCl, 20mM Tris base, pH=7.6 | Made in-house |
| Triton X-100 | 1% TritonX-100 (T9284, Sigma-Aldrich), 150 mM NaCl, 20mM Tris base, 20mM NaF, pH=7.6 | Made in-house |
| urea/SDS | 8M urea, 5% w/v SDS, 150 mM NaCl, 20mM Tris base | Made in-house |
| 4x Laemmli | 0.02% bromophenol blue, 4.4% lithium dodecyl sulphate, 44.4% glycerol, 277.8 mM Tris HCl, pH=6.8 | BioRad (1610747) |
| TBST | 0.1% v/v Tween-20, 150 mM NaCl, 20mM Tris base, pH=7.4 | Made in-house |

Next, we assessed the amount of α-tubulin, a soluble control protein, in each of these fractions. We detected most of the protein in the TBS-soluble fraction (fraction 1), with no significant differences between protocols (Fig. 2B, D). Although some residual α-tubulin was detected in the other two fractions, the amounts were never significant (Fig. 2D). Therefore, α-tubulin remained in the soluble fraction irrespective of the fractionation protocol employed, in contrast with the segregation observed for α-synuclein along the different solubility fractions and protocols.

Furthermore, we employed the same exact protocols for control samples expressing *LacZ* instead of *hSNCA*. In this case the specific 15kDa band that the α*-*synuclein antibody (4B12) detected did not appear in any fractions of head lysates (Fig. 2B, Fig. S1), confirming the specificity of the signal and the lack of α*-*synuclein in flies that do not carry the *UAS-hSNCA* transgene. On the other hand, the soluble control protein, α-tubulin, was detected at a similar level in all the TBS-soluble fractions but remained undetectable or at a very low level in the detergent-soluble or insoluble fractions, regardless of the protocol employed (Fig. S1).

### Sonication increases α-synuclein solubility in SDS buffer

Our results indicated that most α*-*synuclein from 20-day-old fly brains was detected in the detergent-soluble fraction (fraction 2), with minimal amounts in the insoluble (urea) fractions. This contrasts with past literature that reported histological evidence of highly aggregated α*-*synuclein in a similar *Drosophila* model (22), strongly suggesting the presence of insoluble α*-*synuclein. We then questioned whether the protocol that we adapted from the vertebrate literature had been fully optimized for *Drosophila*. In this protocol, we sonicate samples immediately after homogenization. Sonic cavitation produces pressure fluctuations of gigapascal magnitude, generating high-energy mechanical shockwaves that have been reported to degrade some proteins (35). We hypothesized that this high-energy process could be disaggregating physiologically insoluble α*-* synuclein in the protein extracts. To investigate this possibility, we compared the distribution of α*-* synuclein from fly heads that were either sonicated or non-sonicated before biochemical fractionation across the three previous fractions.

We first compared the effects of sonication on α*-*synuclein fractionation under the protocol that employed SDS as the detergent solvent in fraction 2. Under this protocol (Fig. 3A), we did not observe significant differences between sonicated and non-sonicated samples for the TBS-soluble and SDS-soluble fractions (Fig. 3B-C). However, we found a significant increase of about two-fold in the amount of insoluble α*-*synuclein detected in the SDS-insoluble fraction (fraction 3) without sonication when compared to sonicated samples (Fig. 3B-C). We then evaluated the distribution of α*-*tubulin across these three fractions upon either sonication or non-sonication. Regardless of sonication, α*-*tubulin was still predominantly present in the TBS-soluble fraction with almost undetectable traces of the protein in the SDS-soluble and SDS-insoluble fractions (Fig. 3B, D). These data indicated that sonication did not influence cell lysis or protein extraction efficiency as the levels of α*-*tubulin were equivalent for both protocols, sonicated and non-sonicated. In contrast, these results support the conclusion that sonication affects α*-*synuclein solubility, perhaps by breaking down the aggregates. This could explain why sonicated samples herein were found mostly in the SDS-soluble fraction, while non-sonicated samples exhibit higher levels of α*-*synuclein in the SDS-insoluble fraction.

**Figure 3.**
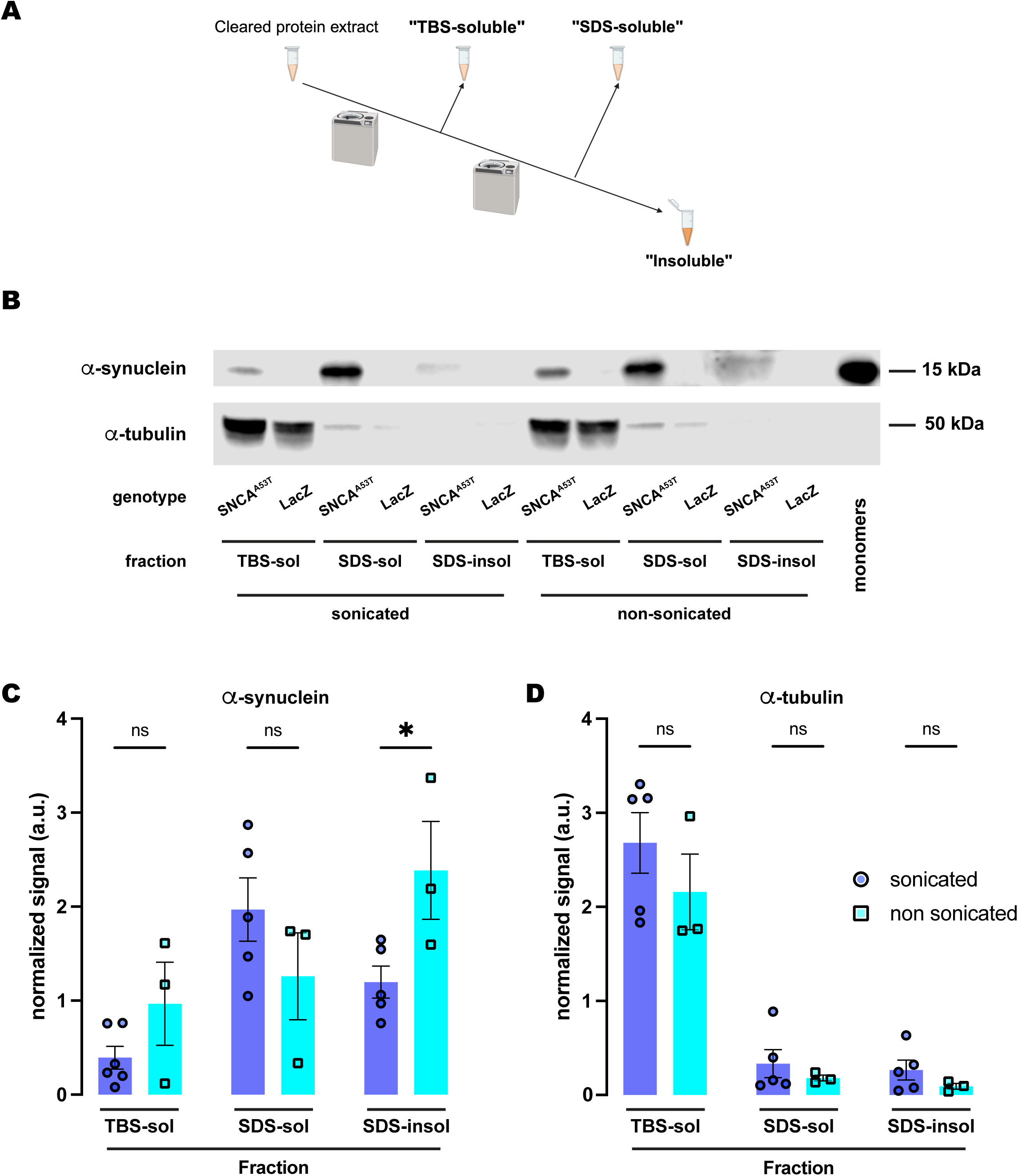
Sonication increases solubility of human A53T α-synuclein in the SDS buffer. **(A)** Schematic representation of the sequential fractionation protocol and the extraction buffers employed in this experiment. **(B)** Representative western blot of head lysates from flies expressing *hSNCA^A53T^* or *LacZ* in dopaminergic neurons. Fly heads are homogenized and then +/- sonication prior to fractionation using a 3-step protocol in which the second fraction uses SDS as the detergent solvent. The first fraction (TBS-soluble) is loaded in lanes 1, 2, 7, 8; the second fraction (SDS-soluble) is loaded in lanes 3, 4, 9, 10; and the third fraction (insoluble) is loaded in lanes 5, 6, 11, 12; while 2ng of purified recombinant human α-synuclein monomers (monomer) are loaded in lane 13 as positive control. Protein lysates are extracted from flies expressing *hSNCA^A53T^*in dopaminergic neurons (*w; +/+; TH-Gal4/UAS-hSNCA^A53T^*, lanes 1, 3, 5, 7, 9, 11) and control flies (*w; +/+; TH-Gal4/UAS-LacZ*) as negative controls not expressing hSNCA (lanes 2, 4, 6, 8, 10, 12). The fractions are probed for α-synuclein (4B12, top panel) and α-tubulin (T6074, bottom panel). **(C)** Quantification of α-synuclein content shows no significant differences between sonicated and non-sonicated samples in the TBS- and SDS-soluble fractions but a significant increase in the amount of insoluble α-synuclein in the third, SDS-insoluble, fraction (Tukey’s multiple comparisons; TBS-soluble, p=0.4969; SDS-soluble, p=0.3459; SDS-insoluble, p=0.0418). **(D)** Quantification of α-tubulin content shows no significant differences between sonication regimens (Tukey’s multiple comparisons; TBS-soluble, p=0.3495; SDS-soluble, p=0.9578; SDS-insoluble, p=0.9398). Samples size was n=5 for sonicated samples and n=3 for non-sonicated samples. Error bars indicate SEM. Tukey’s comparison test: n.s. not significant, * p<0.05, ** p<0.005, *** p<0.001.

### Sonication does not affect α-synuclein solubility in RIPA buffer

We next tested whether sonication elicited a similar effect on the fractionation of α*-*synuclein when the buffer employed was RIPA. Interestingly, the amount of α*-*synuclein in the three fractions (Fig. 4A) was statistically equivalent when comparing sonicated versus non-sonicated samples (Fig. 4B-C). The results revealed that α*-*synuclein was still undetectable in the RIPA-insoluble fraction, while the RIPA-soluble fraction retained most of the protein (Fig. 4B-C). Therefore, we did not observe any changes due to sonication when using RIPA buffer as the buffer in fraction 2 (Fig.4B-C). We then assessed α*-*tubulin and observed similar results to those obtained upon sonication. There were no significant changes between sonicated and non-sonicated samples for any of the fractions, while α*-*tubulin was predominantly present in the TBS-soluble fraction and almost undetectable in the RIPA-soluble and RIPA-insoluble fractions (Fig. 4B, D). Together, these data indicated that sonication has no effect on α*-*synuclein fractionation when using RIPA buffer and suggested a superior extraction efficiency of the detergent solvents contained in this buffer when evaluating the solubility of α*-*synuclein, irrespective of its aggregation state due to sonication.

**Figure 4.**
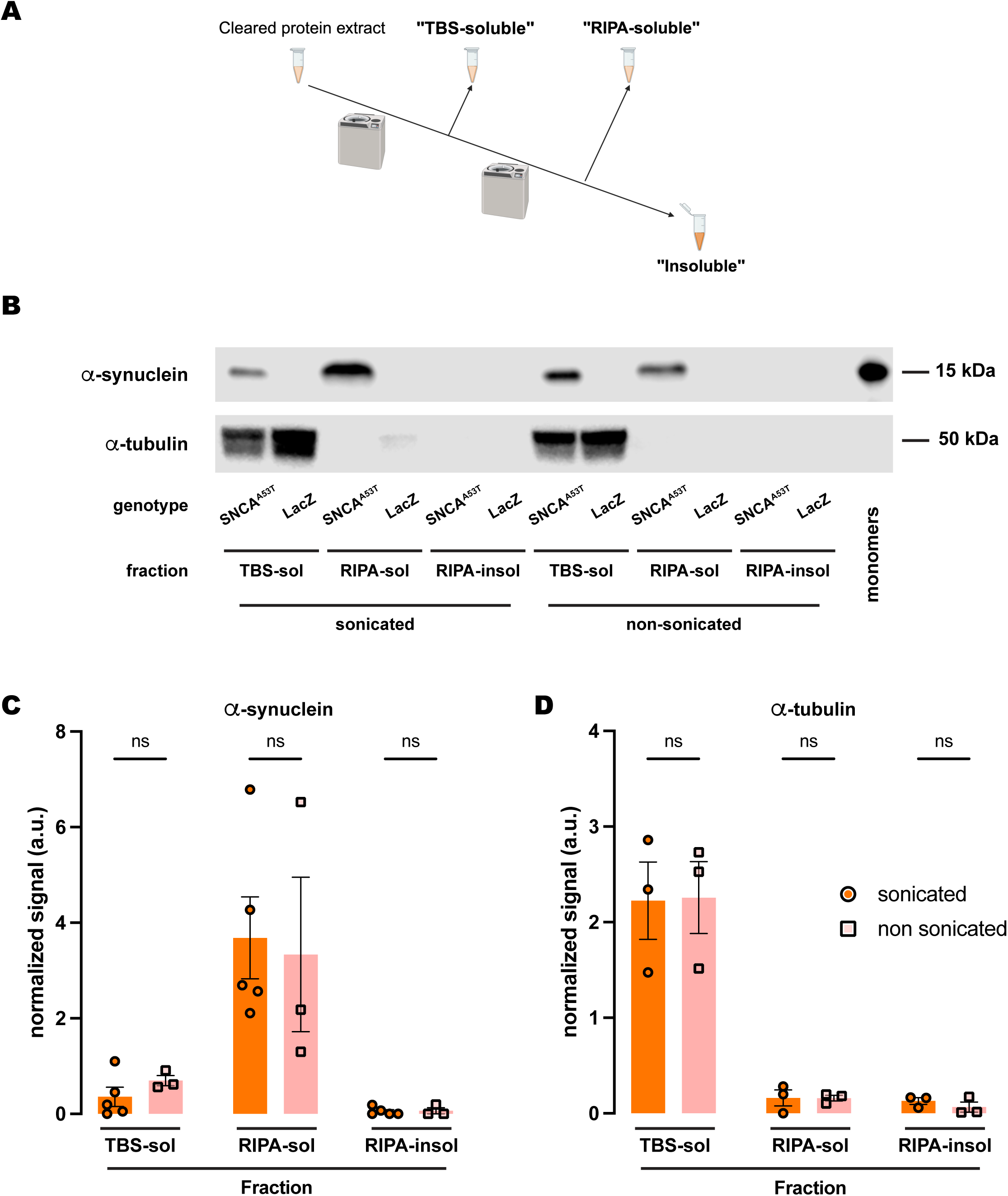
Sonication does not affect human A53T α-synuclein solubility in RIPA buffer. **(A)** Schematic representation of the sequential fractionation protocol and the extraction buffers employed in this experiment. **(B)** Representative western blot of head lysates from flies expressing *hSNCA^A53T^* or *LacZ* in dopaminergic neurons. Fly heads were homogenized and then +/-sonication prior to fractionation using a 3-step protocol in which the second fraction uses RIPA buffer. The first fraction (TBS-soluble) was loaded in lanes 1, 2, 7, 8; the second fraction (RIPA-soluble) was loaded in lanes 3, 4, 9, 10; and the third fraction (insoluble) was loaded in lanes 5, 6, 11, 12; while 2ng of purified recombinant human α-synuclein monomers (monomer) were loaded in lane 13 as positive control. Protein lysates were extracted from flies expressing *hSNCA^A53T^* in dopaminergic neurons (*w; +/+; TH-Gal4/UAS-hSNCA^A53T^*, lanes 1, 3, 5, 7, 9, 11) and control flies (*w; +/+; TH-Gal4/UAS-LacZ*) as negative controls not expressing hSNCA (lanes 2, 4, 6, 8, 10, 12). The fractions were probed for α-synuclein (4B12, top panel) and α-tubulin (T6074, bottom panel). **(C)** Quantification of α-synuclein content shows no significant differences between sonicated and non-sonicated samples in the any of the three fractions (Tukey’s multiple comparisons; TBS-soluble, p=0.9803; RIPA-soluble, p=0.9784; RIPA-insoluble, p>0.9999). α-synuclein is not significantly detected in the RIPA-insoluble fraction (Student’s t-test from zero: sonicated, p=0.2329; non-sonicated, p=0.4226). **(D)** Quantification **of** α-tubulin content shows no significant differences between sonication regimens (Tukey’s multiple comparisons; TBS-soluble, p=0.9995; RIPA-soluble, p>0.9999; RIPA-insoluble, p=0.9965). Samples size was n=5 (α-synuclein) and n=3 (α-tubulin) for sonicated samples, and n=3 for non-sonicated samples. Error bars indicate SEM. Tukey’s comparison test: n.s. not significant, * p<0.05, ** p<0.005, *** p<0.001.

### α*-*synuclein is fully soluble in polyethoxylate detergents irrespective of sonication

Given the discrepancy between the SDS and RIPA buffers regarding α*-*synuclein solubility upon sonication, we decided to further investigate the chemical properties underlying this behavior. Since the RIPA buffer is composed of sodium deoxycholate, NP-40, a polyethoxylated detergent, and sodium dodecyl sulfate (SDS) at a concentration 50X lower than the SDS buffer, we posed the question of whether polyethoxylated detergents could be conferring the differential properties of RIPA versus SDS-alone solubilization. To address this, we performed an experiment using a similar 3-fraction protocol (Fig. 5A) with NP-40 as the detergent solvent of the buffer employed in the second fraction and compared the effects of sonication. Similar to what we observed for the protocol with RIPA buffer, α*-*synuclein was undetectable in the NP-40-insoluble fraction (sonicated and non-sonicated), while the fraction with most α*-*synuclein was the NP-40-soluble fraction (Fig. 5B-C). The amount of α*-*synuclein in each fraction was statistically equivalent regardless of whether or not the samples were previously sonicated (Fig. 5B-C). We also evaluated α*-*tubulin and obtained the exact same results as for RIPA or SDS as detergent solvents (Fig. 3-4), mostly present in the TBS-soluble fraction but undetectable or at a very low level for NP-40-soluble and NP-40-insoluble fractions (Fig. 5B, D). Together, these results lead us to conclude that NP-40, a polyethoxylated detergent, is responsible for the high efficiency with which α*-*synuclein is solubilized in RIPA buffer irrespective of sonication before fractionation.

**Figure 5.**
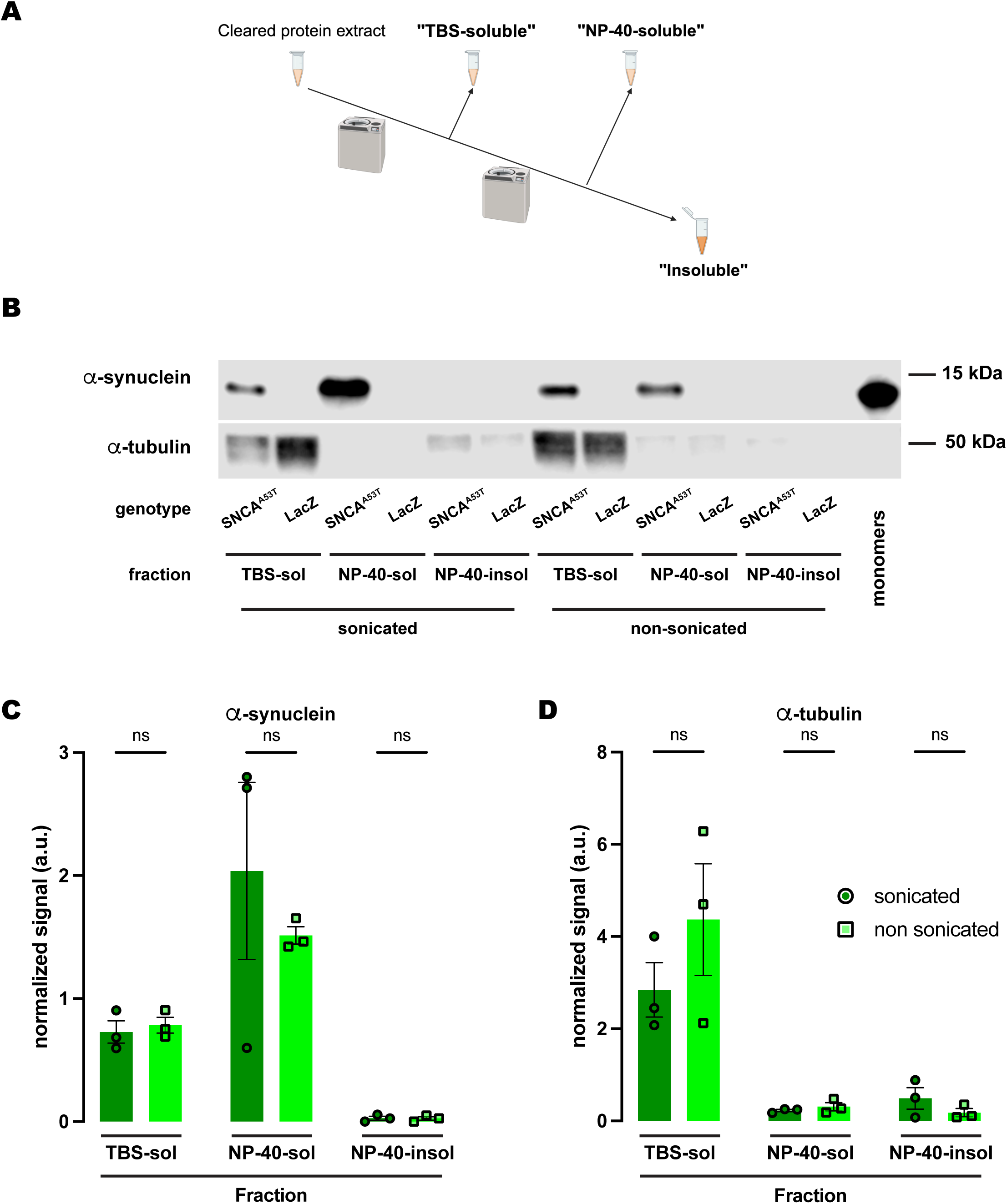
Human A53T α*-*synuclein solubility in buffers containing the polyethoxylate detergent NP-40. **(A)** Schematic representation of the sequential fractionation protocol and the extraction buffers employed in this experiment. **(B)** Representative western blot of head lysates from flies expressing *hSNCA^A53T^* or *LacZ* in dopaminergic neurons. Fly heads are homogenized and then +/- sonication prior to fractionation using a 3-step protocol in which the second fraction uses NP-40 as detergent solvent. The first fraction (TBS-soluble) is loaded in lanes 1, 2, 7, 8; the second fraction (NP-40-soluble) is loaded in lanes 3, 4, 9, 10; the third fraction (insoluble) is loaded in lanes 5, 6, 11, 12; and 2ng of purified recombinant human α-synuclein monomers (monomer) are loaded in lane 13 as positive control. Protein lysates are extracted from flies expressing *hSNCA^A53T^* in dopaminergic neurons (*w; +/+; TH-Gal4/UAS-hSNCA^A53T^*, lanes 1, 3, 5, 7, 9, 11) and control flies (*w; +/+; TH-Gal4/UAS-LacZ*) as negative controls not expressing hSNCA (lanes 2, 4, 6, 8, 10, 12). The fractions are probed for α-synuclein (4B12, top panel) and α-tubulin (T6074, bottom panel). **(C)** Quantification of α-synuclein content shows no significant differences between sonicated and non-sonicated samples in the any of the three fractions (Tukey’s multiple comparisons, sonicated vs non-sonicated: TBS-soluble, p=0.9990; NP-40-soluble, p=0.5587; NP-40-insoluble, p>0.9999). α-synuclein is not significantly detected in the NP-40-insoluble fraction (Student’s t-test from zero: sonicated, p=0.2292; non-sonicated, p=0.2201). **(D**) Quantification of α-tubulin shows with no significant differences between sonication regimens (Tukey’s multiple comparisons, sonicated vs non-sonicated: TBS-soluble, p=0.2177; NP-40-soluble, p=0.9994; NP-40-insoluble, p=0.9741). Samples size was n=3 for sonicated samples and n=3 for non-sonicated samples. Error bars indicate SEM. Tukey’s comparison test: n.s. not significant, * p<0.05, ** p<0.005, *** p<0.001.

Last, we tested whether another polyethoxylated detergent commonly used for fractionation of α*-* synuclein in vertebrate animal models, Triton X-100, conferred the same chemical properties as NP-40 regarding α*-*synuclein solubility. In this case, we employed a 2-step protocol (Fig. 6A), containing Triton X-100-soluble and Triton X-100-insoluble fractions. Similarly, α*-*synuclein was undetectable in the Triton X-100-insoluble fraction regardless of whether samples were previously sonicated, remaining entirely in the X-Triton-soluble fraction for the sonicated and non-sonicated samples (Fig. 6B-C). We also evaluated α*-*tubulin, which was only detected in the Triton X-100-soluble fraction (Fig. 6B, D). In summary, the solubility of α*-*synuclein in polyethoxylated detergents is such that these detergents sequester all α*-*synuclein present in the protein extracts from *Drosophila* heads, regardless of whether they have been previously sonicated or not. In our hands, fractionation buffers containing SDS alone uniquely provide sufficient dynamic range for evaluation of the solubility state of human α*-*synuclein expressed in *Drosophila* brains with high resolution.

**Figure 6.**
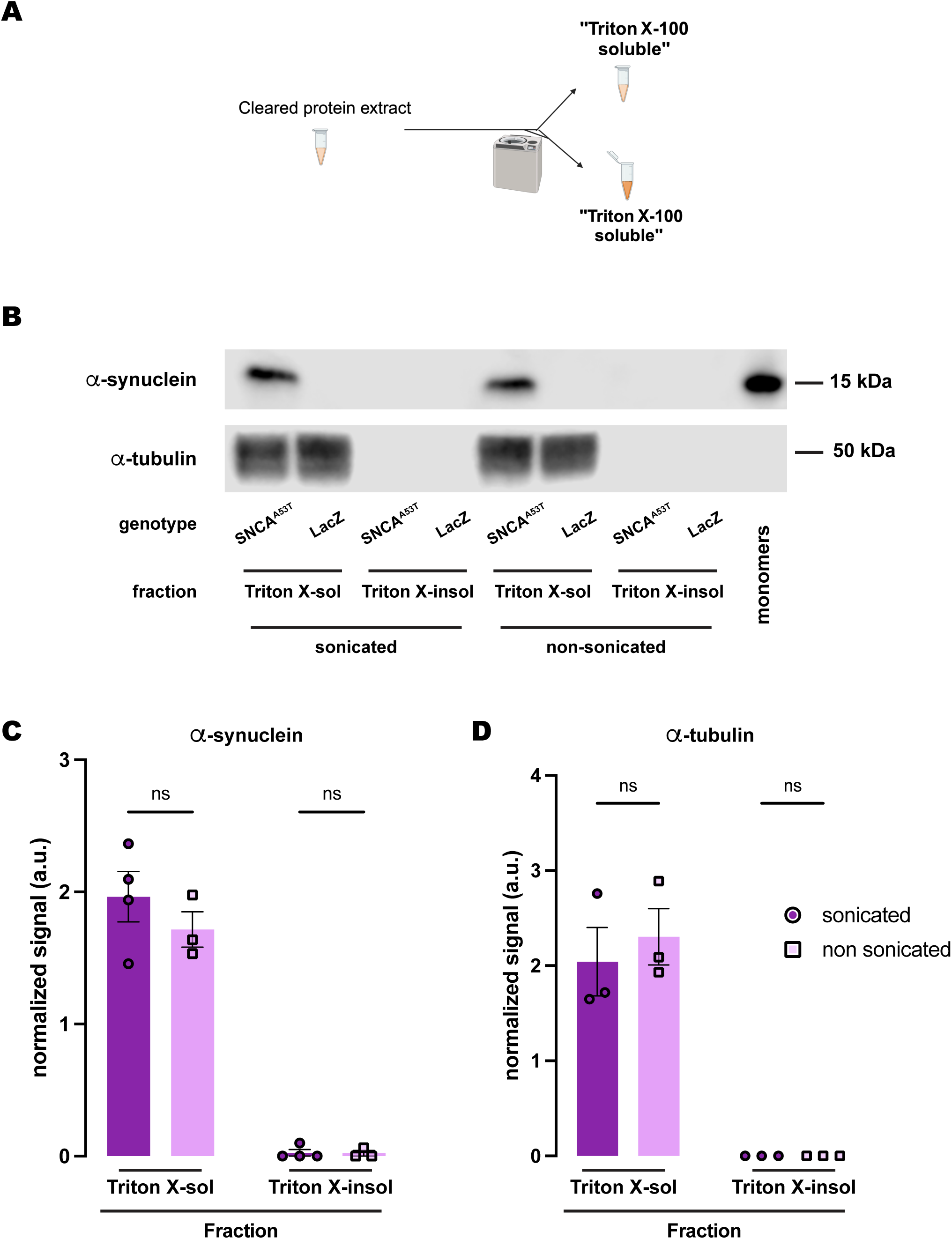
Human A53T α*-*synuclein solubility in buffers containing the polyethoxylate detergent Triton X-100. **(A)** Schematic representation of the sequential fractionation protocol and the extraction buffers employed in this experiment. **(B)** Representative western blot of head lysates from flies expressing *hSNCA^A53T^* or *LacZ* in dopaminergic neurons. Fly heads are homogenized and then +/-sonication prior to fractionation employing a 2-step protocol with Triton X-100 as detergent solvent. The first fraction (Triton X-100-soluble) is loaded in lanes 1, 2, 5, 6; the second fraction (Triton X-100-insoluble) is loaded in lanes 3, 4, 7, 8; and 2ng of purified recombinant human α-synuclein monomers (monomer) are loaded in lane 9 as positive control. Protein lysates are extracted from flies expressing *hSNCA^A53T^*in dopaminergic neurons (*w; +/+; TH-Gal4/UAS-hSNCA^A53T^*, lanes 1, 3, 5, 7) and control flies (*w; +/+; TH-Gal4/UAS-LacZ*) as negative controls not expressing hSNCA (lanes 2, 4, 6, 8). The fractions are probed for α-synuclein (4B12, top panel) and α-tubulin (T6074, bottom panel). **(C-D)** Quantification of α-synuclein **(C)** and α-tubulin **(D)** content shows no significant differences between sonicated and non-sonicated samples in the any of the two fractions (Tukey’s multiple comparisons, sonicated vs non-sonicated: For α-synuclein Triton-X-100-soluble, p=0.3562; Triton-X-100-insoluble, p=0.9998; For α-tubulin Triton-X-100-soluble, p=0.6973; Triton-X-100-insoluble, p>0.9999). α-synuclein is significantly detected in the Triton X-100-soluble fraction (Student’s t-test from zero: sonicated, p=0.0019; non-sonicated, p=0.0061) but not significantly in the Triton-X-100-insoluble fraction (Student’s t-test from zero: sonicated, p=0.3910; non-sonicated, p=0.4226). α-tubulin is significantly detected in the Triton X-100-soluble fraction (Student’s t-test from zero: sonicated, p=0.0296; non-sonicated, p=0.0162) and not detected in the Triton-X-100-insoluble fraction. Samples size was n=4 (α-synuclein) and n=3 (α-tubulin) for sonicated samples, and n=3 for non-sonicated samples. Error bars indicate SEM. Tukey’s comparison test: n.s. not significant, * p<0.05, ** p<0.005, *** p<0.001.

## DISCUSSION

Chemical fractionation of misfolded and aggregated proteins is a critical method to assess the solubility and the species composition of the proteinaceous deposits formed by these proteins and their relationship to disease stage. We tested a variety of protocols to evaluate the solubility of *hSNCA* expressed in DA neurons of the *Drosophila* brain and found that buffers containing polyethoxylated detergents such as NP-40 or Triton X-100 exert such a strong extraction efficiency for TBS-insoluble α-synuclein that no α-synuclein remains in the insoluble fraction. However, SDS-containing solvents only solubilize a fraction of the remaining α-synuclein, leaving the most insoluble molecules in the precipitate. We also observed that sonication prior to chemical fractionation breaks down α-synuclein insoluble aggregated species, making them soluble in the SDS-containing fraction. Below, we consider several possible explanations for this biochemical behavior of human α-synuclein.

First, we looked at the chemical properties of these detergent solvents seeking to find the basis for the difference in extraction efficiency. SDS is an anionic detergent that consists of a short acyl chain attached to a sulfate head group. On the other hand, RIPA is a composite detergent buffer that contains 1% NP-40 (a nonionic polyethoxylate detergent with 24-51 ethyl ether groups per molecule), 1% sodium deoxycholate (an anionic detergent that consists of a steroid core attached to a carboxylate head group), and 0.1% SDS. At these concentrations, all these molecules form micelles (critical micelle concentrations: SDS - .079% w/v (36), NP-40 - .035% w/v (37), sodium deoxycholate - .099% w/v (38) that encapsulate the hydrophobic portions of proteins to various degrees, extracting them into solution. We suspect that the presence of NP-40 is responsible for the higher content of α-synuclein in the RIPA-soluble fractions, given that the use of NP-40 alone mimicked the results obtained with RIPA. With 24-51 ethyl ether groups (37), NP-40 molecules can be 87-168 Å long, which is 7-14 times longer than an SDS molecule (Fig. 7). Moreover, the average number of molecules per micelle (as measured by the aggregation number, N_A_) for NP-40 is around 150 (39), more than twice as large as that of SDS [N_A_=67; (40)]. As a result of these biophysical properties, NP-40 forms larger and more flexible micelles, which we hypothesize are more capable of encapsulating and extracting larger α-synuclein complexes than the smaller SDS micelles. Therefore, RIPA buffer as a solubilization solvent may lead to an underestimation of the insoluble human α-synuclein content due to its higher capacity to sequester larger α-synuclein insoluble aggregates. We tested this hypothesis by extracting α-synuclein using a buffer with 1% NP-40 as detergent and found that NP-40-soluble fractions were highly enriched in α-synuclein, while α-synuclein was almost undetectable in NP-40-insoluble fraction. These data recapitulate the results we observed when using RIPA, consistent with our hypothesis.

**Figure 7.**
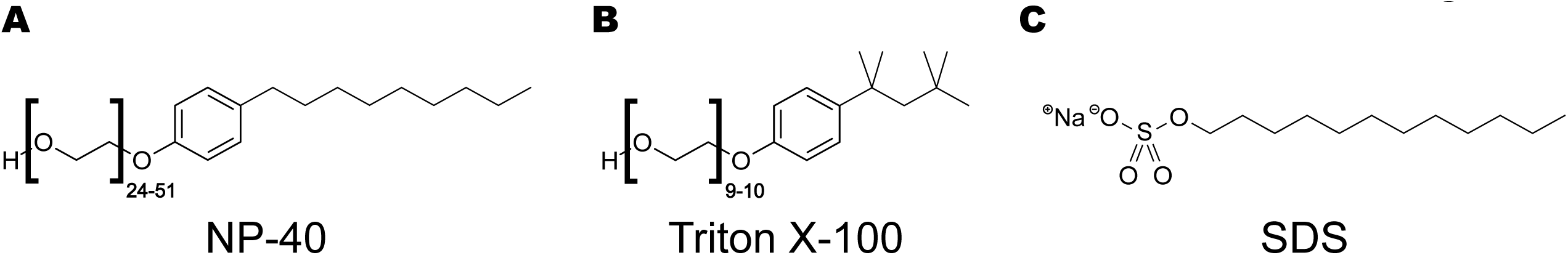
Chemical structures of the detergents NP-40, Triton X-100 and SDS employed in this study. **(A)** NP-40, polyethylene glycol nonyl-phenyl ether or nonoxynol-40. **(B)** Triton X-100, polyethylene glycol *p*-(1,1,3,3-tetramethylbutyl)-phenyl ether, or octyl phenol ethoxylate. **(C)** Sodium dodecyl sulfate (SDS). Note the structural similarities between the polyethoxylated detergents, NP-40 and Triton X-100, with long chains of ethyl ether groups (24-51 for NP-40 and 9-10 for Triton X-100), which are lacking from the structure of SDS.

Another detergent that is commonly used in the field to assess insoluble α-synuclein levels is Triton X-100, a nonionic polyethoxylate detergent very similar to NP-40. Earls et al. (2020) used suspensions of Triton X-100-insoluble material to measure the uptake of aggregated α-synuclein in human NK cell cultures from media containing pre-formed fibrils (PFFs). In an independent study, Quintin et al. used the same approach to assess α-synuclein aggregation in HEK cells incubated with PFFs (6). To the best of our knowledge, Triton X-100 is the only detergent used for α-synuclein biochemical fractionation in the *Drosophila* literature. Miura et al., Suzuki et al., Davis et al., and Khair et al. (17, 19, 26, 27) used Triton X-100-insoluble α-synuclein as a measure of protein aggregation. Suzuki et al. obtained a very weak α-synuclein monomer signal in the insoluble fraction, many times weaker than soluble α-synuclein signal, with no higher molecular weight insoluble α-synuclein species detected(27). Miura et al. did not detect any insoluble α-synuclein monomers but reported detecting a weak α-synuclein signal at 140kDa(26). Similarly, Davis et al. also reported the ability to detect soluble but not insoluble α-synuclein monomers while they observe higher molecular weight species in the insoluble fraction(19). We were unable to detect Triton X-100-insoluble α-synuclein. The undetectability of Triton X-100-insoluble α-synuclein in our expression system is internally consistent with the rest of our experimental findings herein. As mentioned earlier, Triton X-100 is a polymer detergent with repeating ethyl ether groups. It differs from NP-40 only in the alkyl group attached to the polyethoxylated phenyl ring (tetramethylbutyl in TritonX and nonyl in NP-40) and a fewer number of ethyl ether repeats. Therefore, we would expect Triton X-100 to form micelles similar in their physical and chemical properties to those formed by NP-40, thus extracting α-synuclein with a similar efficiency. The undetectability of detergent-insoluble α-synuclein in RIPA buffer, NP-40-, and Triton X-100- containing solvents suggests that the repeating ethyl ether groups are responsible for the higher solubilization of α-synuclein in buffers containing these detergents.

One potential reason for not detecting detergent-insoluble α-synuclein, which differs from previously published data, may be the lower overall expression of human α-synuclein in our model; the aforementioned *Drosophila* studies expressed human α-synuclein to higher levels, either pan-neuronally with *elav* or ubiquitously with actin promoters, while we intentionally restrict expression to DA neurons (∼130 per protocerebrum hemisphere), which constitutes only a small percentage of all neurons in the fly brain (<200,000 neurons). It is likely that α-synuclein is not processed in the same way in every neuronal type and, therefore, we focus our study on a more physiologically relevant neuronal population, DA neurons, widely known to develop α-synuclein pathology and degenerate in PD. In addition, it is possible that our driver is not only more selective, but it may also drive weaker expression than other ubiquitous promoters such as that from actin. Relative abundance of α-synuclein could both increase the aggregation rate, as mutations that lead to higher expression levels of α-synuclein trigger aggregation and LB pathology (12–14), with overall higher total accumulation of aggregated α-synuclein leading to more robust and resistant detection of aggregated α-synuclein via standard biochemical fractionation and detection methods.

An additional critical factor to keep in mind when performing these fractionation experiments is whether to perform sample sonication before the serial fractionation. In our experiments using SDS as the detergent solvent in the fractionation buffer, sonication of fly head homogenates before biochemical fractionation induced enrichment in α-synuclein content in the SDS-soluble fraction while reducing the amount of total α-synuclein in the SDS-insoluble fraction. These findings demonstrate that sonication is sufficient to solubilize physiologically insoluble α-synuclein accumulated in *Drosophila* DA neurons and suggest that, to a certain degree, sonication triggers disaggregation of α-synuclein complexes. For this reason, we would recommend that, when using *Drosophila* to model synucleinopathies, sonication before chemical fractionation should be avoided to prevent confounding interpretation of its solubility state *in vivo*. Contrary to the need for sonication in samples from vertebrate animals that are rich in myelin and other fatty components, fly samples do not require sonication to break down those additional components present in vertebrates. Therefore, the use of sonication prior to serial fractionation is unnecessary and likely undesirable as it will compromise the interpretation of the biophysical and biochemical properties of the α-synuclein species that are being extracted and studied.

Regardless of protocol, we could not detect solubilized α-synuclein of a higher molecular weight than that of recombinant α-synuclein monomeric protein. It is important to note that westerns were not conducted under complete native conditions. However, protein lysate samples were not reduced with BME or boiled, yet they were diluted with SDS-containing Laemmli buffer prior to loading in the native gel, which likely had an effect on the protein’s natural structure and may account for the monomeric weight. Nevertheless, the limitation of the model in detecting higher molecular weight species is likely due to the low amount of α-synuclein that we find in the brain of these flies, as discussed earlier. However, it is also possible that *Drosophila* DA neurons expressing *hSNCA* do not produce high molecular weight species as found in LBs, which typically require tough solvents such as urea or acid to solubilize the large α-synuclein aggregates (41, 42).

Importantly, the choice of detergent for biochemical fractionation depends on the research question and the conditions suitable for the model. In cases when the detergent-soluble protein content is used as a measure of membrane-associated protein, our data demonstrate that SDS is a better detergent for these purposes. SDS, being a fatty acid similar in length to a typical phospholipid, resembles the plasma membrane more than RIPA (NP-40) or Triton X-100 do, because they contain considerably longer polymer detergents. In addition, extraction with SDS allows a more direct comparison to the existing literature in human postmortem brains when assessing α-synuclein solubility, which has mainly employed SDS as a detergent in the serial biochemical fractionation of α-synuclein (15, 16, 43–45). In conclusion, when selecting an appropriate fractionation protocol for Drosophila studies, our findings support the usage of a 3-step fractionation protocol where the buffer for the second fraction contains only SDS at 5% concentration. The combination of a buffer with these biochemical properties and the multi-fraction protocol provides a dynamic range capable of resolving several degrees of insolubility in human α-synuclein processed in DA neurons of the *Drosophila* brain.

## EXPERIMENTAL PROCEDURES

### Fly strains

Fly stocks were raised on standard corn syrup-soy (W1) food (Bloomington stock center recipe) from Archon Scientific. All crosses were maintained at 25°C and 70% relative humidity on a 12 hours light:dark cycle. Flies carrying the *TH-Gal4* driver were provided by Dr. Matthew Kayser from University of Pennsylvania. Expression lines UAS-LacZ (BDSC, catalog #8530; RRID:BDSC_8530) and UAS-hSNCA^A53T^ (BDSC, catalog #8148; RRID:BDSC_8148) were obtained from the Bloomington Stock center. Line for expression of the human α-synuclein bearing the pathology-associated mutation A53T (hSNCA^A53T^) was previously described (22). Flies were collected immediately upon eclosion and housed in individual vials in groups of 20 males. All experiments were performed in flies aged to 20 days post-eclosion.

### Tissue collection and fractionation

#### 3-step fractionation protocol

Groups of 20 male flies were transferred to Eppendorf cryovials, snap-frozen in liquid nitrogen and then vortexed. 20-40 fly heads were then collected into BioMasher® II Tissue Grinder (DWK Life Sciences, 749625-0020) tubes and homogenized for 1.5 minutes in 50μL of Tris-buffered saline (TBS; Table 1) with 1X protease and 1X phosphatase inhibitor cocktails (Roche, 04693124001 and 04906837001). The homogenates were then probe-sonicated with a Branson Sonifier® SFX150 (Emerson) at 35% amplitude (∼52.5 watts) for five 1-second pulses, unless otherwise specified. The suspension was clarified by 15-second centrifugation at 2,100*g* and the supernatant was ultracentrifuged at 100,000*g* for 30 minutes at 4°C using the Sorvall WX80 Plua Ultracentrifuge (ThermoScientific). The supernatant was labelled as the TBS-soluble fraction (fraction1). To ensure complete removal of soluble material, the pellet was washed in TBS and ultracentrifuged again at 100,000*g* for 15 min at 4°C, after which the supernatant was discarded. The pellet was then re-homogenized in 30μL of either SDS, NP-40, RIPA, or TBS buffer (Table 1) with protease and phosphatase inhibitors and ultracentrifuged at 100,000*g* for 30 minutes at 4°C. The supernatant (fraction 2) was collected and labelled as SDS-soluble, NP-40-soluble, RIPA-soluble, or TBS-wash, respectively. The remaining pellet was washed in the previous solvent and centrifuged again at 100,000*g* for 15 min at 4°C, after which the supernatant was discarded. The final pellet was resuspended in 30μL urea/SDS (Table 1) with protease and phosphatase inhibitors by probe-sonication with three 1-second pulses at 35% amplitude and the suspension was labelled as the insoluble fraction (fraction 3; Fig. 1A).

#### 2-step fractionation protocol

Groups of 20 male flies were transferred to Eppendorf cryovials, snap-frozen in liquid nitrogen and then vortexed. 20-40 fly heads were then collected into BioMasher® II Tissue Grinder (DWK Life Sciences, 749625-0020) tubes and homogenized for 1.5 minutes in 50μL of TBS containing 1% Triton X-100 (Table 1) and 1X protease and 1X phosphatase inhibitor cocktails (Roche, 04693124001 and 04906837001). The homogenates were then probe-sonicated with a Branson Sonifier® SFX150 (Emerson) at 35% amplitude (∼52.5 watts) for five 1-second pulses, unless otherwise specified. The suspension was clarified by 15-second centrifugation at 2,100*g* and the supernatant was ultracentrifuged at 100,000*g* for 30 minutes at 4°C using the Sorvall WX80 Plua Ultracentrifuge (ThermoScientific). The supernatant was labelled as the Triton X-100-soluble fraction (fraction1). To ensure complete removal of soluble material, the pellet was washed in the Triton X-100 buffer (Table 1) and centrifuged again at 100,000*g* for 15 min at 4°C, after which the supernatant was discarded. The final pellet was resuspended in 50μL of the Triton X-100 buffer (Table 1) with protease and phosphatase inhibitors by probe-sonication with three 1-second pulses at 35% amplitude and the suspension was labelled as the Triton-X-100-insoluble fraction (fraction2; Fig. 1B).

### Quantitative western immunoblotting

The total protein concentration in each fraction was determined by bicinchoninic acid assay (BCA) (Thermo Fisher Scientific, 23225). Then, 20μg of total protein per fraction were collected and diluted with their respective solvents and 4X Laemmli buffer (Biorad, 1610747) to obtain aliquots of equal volume. These samples were loaded into the 10% Bis-Tris polyacrylamide gel (BioRad, 3450112) and subjected to electrophoresis at 80V for 15 minutes and 100V for 90 minutes. After separation, protein was transferred via a semi-wet electrophoretic transfer (Turbo-blot turbo transfer system, Bio-Rad) onto 0.2μm polyvinylidene fluoride (PVDF) membrane membranes (BioRad, 1704273). The PVDF membranes were fixed with 0.8% paraformaldehyde for 30 minutes, washed with H_2_0, and then stained with Revert Total Protein Stain (LI-COR, 926-11016) and the signal used for signal normalization. Membranes were then destained and blocked with 5% powdered milk in TBST buffer, following an overnight incubation at 4°C with the primary antibody against residues 103-108 of human α-synuclein (BioLegend, 807801 [4B12] 1:1000; RRID: AB_2564730). Next day, HRP-conjugated secondary antibody (Jackson Immuno Research, 115-036-072; RRID: AB_2338525) was incubated for 1 hour at room temperature, after which the protein bands were visualized using enhanced chemiluminescence (Thermo Fisher Scientific, 34096) on a LI-COR Odyssey® Fc. The membranes were then stripped from secondary antibody with 0.2M NaOH, re-blocked, and re-probed with the anti-tubulin primary antibody (Sigma-Aldrich, T6074 1:200,000; RRID: AB_477582) and the HRP-conjugated secondary antibody after which they were re-imaged.

### Experimental design and statistical analyses

For all experiments, controls and genetically matched experimental genotypes were performed in parallel. The experimental design ensured that all groups were balanced throughout the experiments conducted. Statistical significance was assessed using GraphPad Prism v10.1 (RRID:SCR_002798). Control flies were balanced for the presence of pUAS constructs by expressing the UAS-LacZ in place of the UAS-hSNCA^A53T^ of the experimental groups. Across all gels, a previously tested sample of known antibody reactivity was run as the internal control sample that enabled comparison across blots. The raw protein band signal was first normalized to total protein staining to control for loading differences and then reported as the fold difference from the internal control sample signal. Ordinary two-way ANOVA test followed by Tukey’s multiple *post-hoc* comparisons were employed to analyze statistical significance across fractions and solvents. Student’s t-test was used to assess significance from zero and determine whether there were significant levels of α-synuclein in the fraction. All data presented represent mean ± the SEM.

## DATA AVAILABILITY

The authors confirm that the data supporting the findings of this study are available within the article and its supporting information. Raw data will be available upon request from the corresponding author.

## SUPPORTING INFORMATION

This article contains supporting information.

## ACKNOWLEDGEMENTS

Partial funding for this work was derived from awards from the Alzheimer’s Association AARG-D-22-972117 (AMP), the Center for Translational Research in Neurodegenerative Disease (AMP), the Parkinson’s Foundation (AMP/MGT), NIH NINDS 1RF1NS28800 (MGT), and the joint efforts of The Michael J. Fox Foundation for Parkinson’s Research (MJFF) and the Aligning Science Across Parkinson’s (ASAP) initiative. MJFF administers the grant ASAP-020527 on behalf of ASAP and itself. We thank Dr. Benoit Giasson from the University of Florida for extensive discussions about the different fractionation protocols and solvents. Stocks obtained from the Bloomington Drosophila Stock Center (NIH P40OD018537) were used in this study. The content of this manuscript is solely the responsibility of the authors and does not necessarily represent the official views of the National Institute of Health.

---

**Figure S1 (Related to figure 2). Lack of antibody reactivity in genetic control flies. (A)** Schematic representation of the sequential fractionation protocol and the extraction buffers employed in this experiment. **(B)** Representative western blot of head lysates from flies expressing *LacZ* or *hSNCA^A53T^* in dopaminergic neurons. Fly heads are fractionated using a 3-step protocol in which the second fraction uses a variable detergent solvent, TBS, SDS or RIPA buffer. The first fraction (TBS-soluble) is loaded in lanes 1-4, second fraction (TBS-wash, SDSsoluble or RIPA-soluble) is loaded in lanes 5-8, and the third fraction (insoluble) is loaded in lanes 9-12, while 2ng of purified recombinant human α-synuclein monomers (monomer) are loaded in lane 13 as positive control. Protein lysates are extracted from flies expressing *LacZ* in dopaminergic neurons (*w*; +/+; *TH-Gal4/ UAS-LacZ*, lanes 1-3, 5-7, 9-11) and flies expressing *hSNCA^A53T^* (*w*; +/+; *TH-Gal4/ UAS-hSNCA^A53T^*, lanes 4, 8, 12). The fractions are probed for α- synuclein (4B12, top panel) and α-tubulin (T6074, bottom panel). Note that control flies (*w*; +/+; *TH-Gal4/ UAS-LacZ*) do not show any reactivity to the α-synuclein antibody regardless of the fractionation protocol. **(C)** Total protein staining with Revert Total Protein Stain of the membrane employed in this experiment.

**Figure S2 (Related to figure 2).** Total protein staining with Revert Total Protein Stain of the membrane employed for the experiment in figure 2.

**Figure S3 (Related to figure 3).** Total protein staining with Revert Total Protein Stain of the membrane employed for the experiment in figure 3.

**Figure S4 (Related to figure 4).** Total protein staining with Revert Total Protein Stain of the membrane employed for the experiment in figure 4.

**Figure S5 (Related to figure 5).** Total protein staining with Revert Total Protein Stain of the membrane employed for the experiment in figure 5.

**Figure S6 (Related to figure 6).** Total protein staining with Revert Total Protein Stain of the membrane employed for the experiment in figure 6.

**Table S1.** Comparison of α-synuclein enrichment in different fractions. The data show differences in mean values ± SEM of normalized α-synuclein signal detected in each biochemical fraction from *TH-Gal4/UAS-hSNCA^A53T^* flies (sonicated and non-sonicated data were grouped for this analysis), and p-values from *post-hoc* Tukey’s multiple comparisons assessing the significance of the differences between the compared groups. Normalization was performed in reference to total protein staining. Signal values are in arbitrary units (a.u.).

| **Fractionation Sequence** | **Fractions compared (fraction A – fraction B)** | **Difference (fraction A – fraction B)** | **p-value** |
| --- | --- | --- | --- |
| **TBS>SDS>insol** | TBS - SDS | -1.119 ± 0.353 | 0.0120 |
| | TBS - Insol | -1.059 ± 0.353 | 0.0176 |
| | SDS - Insol | 0.060 ± 0.364 | 0.9849 |
| **TBS>RIPA>insol** | TBS - RIPA | -3.068 ± 0.614 | 0.0002 |
| | TBS - Insol | 0.429 ± 0.614 | 0.7676 |
| | RIPA - Insol | 3.496 ± 0.614 | <0.0001 |
| **TBS>NP-40>insol** | TBS - NP-40 | -1.019 ± 0.284 | 0.0071 |
| | TBS - Insol | 0.730 ± 0.284 | 0.0527 |
| | NP-40 - Insol | 1.749 ± 0.284 | <0.0001 |
| **Triton>insol** | TritonX - Insol | 1.834 ± 0.1253 | <0.0001 |

**Table S2.** Comparison of α-tubulin enrichment in different fractions. The data show differences in mean values ± SEM of normalized α-tubulin signal detected in each biochemical fraction from *TH-Gal4/UAS-hSNCA^A53T^* flies (sonicated and non-sonicated data were grouped for this analysis), and p-values from *post-hoc* Tukey’s multiple comparisons assessing the significance of the differences between the compared groups. Normalization was performed in reference to total protein staining. Signal values are in arbitrary units (a.u.).

| **Fractionation Sequence** | **Fractions compared (fraction A – fraction B)** | **Difference (fraction A – fraction B)** | **p-value** |
| --- | --- | --- | --- |
| **TBS>SDS>insol** | TBS - SDS | 2.209 ± 0.227 | <0.0001 |
| | TBS - Insol | 2.284 ± 0.227 | <0.0001 |
| | SDS - Insol | 0.075 ± 0.227 | 0.9418 |
| **TBS>RIPA>insol** | TBS - RIPA | 2.081 ± 0.206 | <0.0001 |
| | TBS - Insol | 2.144 ± 0.206 | <0.0001 |
| | RIPA - Insol | 0.063 ± 0.206 | 0.9493 |
| **TBS>NP-40>insol** | TBS - NP-40 | 3.338 ± 0.577 | 0.0001 |
| | TBS - Insol | 3.271 ± 0.577 | 0.0001 |
| | NP-40 - Insol | -0.067 ± 0.577 | 0.9925 |
| **Triton>insol** | TritonX - Insol | 2.173 ± 0.216 | <0.0001 |

## REFERENCES

1. Goedert, M., and Spillantini, M. G. (1998) Lewy body diseases and multiple system atrophy as alpha-synucleinopathies. Mol Psychiatry. 3, 462–465

2. Tong, J., Wong, H., Guttman, M., Ang, L. C., Forno, L. S., Shimadzu, M., Rajput, A. H., Muenter, M. D., Kish, S. J., Hornykiewicz, O., and Furukawa, Y. (2010) Brain alpha-synuclein accumulation in multiple system atrophy, Parkinson’s disease and progressive supranuclear palsy: a comparative investigation. Brain. 133, 172–188

3. Parra-Rivas, L. A., Madhivanan, K., Aulston, B. D., Wang, L., Prakashchand, D. D., Boyer, N. P., Saia-Cereda, V. M., Branes-Guerrero, K., Pizzo, D. P., Bagchi, P., Sundar, V. S., Tang, Y., Das, U., Scott, D. A., Rangamani, P., Ogawa, Y., and Subhojit Roy, null (2023) Serine-129 phosphorylation of α-synuclein is an activity-dependent trigger for physiologic protein-protein interactions and synaptic function. Neuron. 111, 4006–4023.e10

4. Marotta, N. P., Ara, J., Uemura, N., Lougee, M. G., Meymand, E. S., Zhang, B., Petersson, E. J., Trojanowski, J. Q., and Lee, V. M.-Y. (2021) Alpha-synuclein from patient Lewy bodies exhibits distinct pathological activity that can be propagated in vitro. Acta Neuropathol Commun. 9, 188

5. Ma, L., Yang, C., Zhang, X., Li, Y., Wang, S., Zheng, L., and Huang, K. (2018) C-terminal truncation exacerbates the aggregation and cytotoxicity of α-Synuclein: A vicious cycle in Parkinson’s disease. Biochim Biophys Acta Mol Basis Dis. 1864, 3714–3725

6. Quintin, S., Lloyd, G. M., Paterno, G., Xia, Y., Sorrentino, Z., Bell, B. M., Gorion, K.-M., Lee, E. B., Prokop, S., and Giasson, B. I. (2023) Cellular processing of α-synuclein fibrils results in distinct physiological C-terminal truncations with a major cleavage site at residue Glu 114. J Biol Chem. 299, 104912

7. Moon, S. P., Balana, A. T., Galesic, A., Rakshit, A., and Pratt, M. R. (2020) Ubiquitination Can Change the Structure of the α-Synuclein Amyloid Fiber in a Site Selective Fashion. J Org Chem. 85, 1548–1555

8. Burai, R., Ait-Bouziad, N., Chiki, A., and Lashuel, H. A. (2015) Elucidating the Role of Site-Specific Nitration of α-Synuclein in the Pathogenesis of Parkinson’s Disease via Protein Semisynthesis and Mutagenesis. J Am Chem Soc. 137, 5041–5052

9. Schreurs, S., Gerard, M., Derua, R., Waelkens, E., Taymans, J.-M., Baekelandt, V., and Engelborghs, Y. (2014) In vitro phosphorylation does not influence the aggregation kinetics of WT α-synuclein in contrast to its phosphorylation mutants. Int J Mol Sci. 15, 1040–1067

10. Karampetsou, M., Ardah, M. T., Semitekolou, M., Polissidis, A., Samiotaki, M., Kalomoiri, M., Majbour, N., Xanthou, G., El-Agnaf, O. M. A., and Vekrellis, K. (2017) Phosphorylated exogenous alpha-synuclein fibrils exacerbate pathology and induce neuronal dysfunction in mice. Sci Rep. 7, 16533

11. Moors, T. E., Maat, C. A., Niedieker, D., Mona, D., Petersen, D., Timmermans-Huisman, E., Kole, J., El-Mashtoly, S. F., Spycher, L., Zago, W., Barbour, R., Mundigl, O., Kaluza, K., Huber, S., Hug, M. N., Kremer, T., Ritter, M., Dziadek, S., Geurts, J. J. G., Gerwert, K., Britschgi, M., and van de Berg, W. D. J. (2021) The subcellular arrangement of alpha-synuclein proteoforms in the Parkinson’s disease brain as revealed by multicolor STED microscopy. Acta Neuropathol. 142, 423–448

12. Singleton, A. B., Farrer, M., Johnson, J., Singleton, A., Hague, S., Kachergus, J., Hulihan, M., Peuralinna, T., Dutra, A., Nussbaum, R., Lincoln, S., Crawley, A., Hanson, M., Maraganore, D., Adler, C., Cookson, M. R., Muenter, M., Baptista, M., Miller, D., Blancato, J., Hardy, J., and Gwinn-Hardy, K. (2003) alpha-Synuclein locus triplication causes Parkinson’s disease. Science. 302, 841

13. Chartier-Harlin, M.-C., Kachergus, J., Roumier, C., Mouroux, V., Douay, X., Lincoln, S., Levecque, C., Larvor, L., Andrieux, J., Hulihan, M., Waucquier, N., Defebvre, L., Amouyel, P., Farrer, M., and Destée, A. (2004) Alpha-synuclein locus duplication as a cause of familial Parkinson’s disease. Lancet. 364, 1167–1169

14. Konno, T., Ross, O. A., Puschmann, A., Dickson, D. W., and Wszolek, Z. K. (2016) Autosomal dominant Parkinson’s disease caused by SNCA duplications. Parkinsonism Relat Disord. 22 Suppl 1, S1–6

15. Zhou, J., Broe, M., Huang, Y., Anderson, J. P., Gai, W.-P., Milward, E. A., Porritt, M., Howells, D., Hughes, A. J., Wang, X., and Halliday, G. M. (2011) Changes in the solubility and phosphorylation of α-synuclein over the course of Parkinson’s disease. Acta Neuropathol. 121, 695–704

16. Mamais, A., Raja, M., Manzoni, C., Dihanich, S., Lees, A., Moore, D., Lewis, P. A., and Bandopadhyay, R. (2013) Divergent α-synuclein solubility and aggregation properties in G2019S LRRK2 Parkinson’s disease brains with Lewy Body pathology compared to idiopathic cases. Neurobiol Dis. 58, 183–190

17. Abul Khair, S. B., Dhanushkodi, N. R., Ardah, M. T., Chen, W., Yang, Y., and Haque, M. E. (2018) Silencing of Glucocerebrosidase Gene in Drosophila Enhances the Aggregation of Parkinson’s Disease Associated α-Synuclein Mutant A53T and Affects Locomotor Activity. Front Neurosci. 12, 81

18. Auluck, P. K., Chan, H. Y. E., Trojanowski, J. Q., Lee, V. M. Y., and Bonini, N. M. (2002) Chaperone suppression of alpha-synuclein toxicity in a Drosophila model for Parkinson’s disease. Science. 295, 865–868

19. Davis, M. Y., Trinh, K., Thomas, R. E., Yu, S., Germanos, A. A., Whitley, B. N., Sardi, S. P., Montine, T. J., and Pallanck, L. J. (2016) Glucocerebrosidase Deficiency in Drosophila Results in α-Synuclein-Independent Protein Aggregation and Neurodegeneration. PLoS Genet. 12, e1005944

20. Dhanushkodi, N. R., Abul Khair, S. B., Ardah, M. T., and Haque, M. E. (2023) ATP13A2 Gene Silencing in Drosophila Affects Autophagic Degradation of A53T Mutant α-Synuclein. Int J Mol Sci. 24, 1775

21. Du, G., Liu, X., Chen, X., Song, M., Yan, Y., Jiao, R., and Wang, C.-C. (2010) Drosophila histone deacetylase 6 protects dopaminergic neurons against {alpha}-synuclein toxicity by promoting inclusion formation. Mol Biol Cell. 21, 2128–2137

22. Feany, M. B., and Bender, W. W. (2000) A Drosophila model of Parkinson’s disease. Nature. 404, 394–398

23. Golomidov, I., Bolshakova, O., Komissarov, A., Sharoyko, V., Slepneva, Е., Slobodina, A., Latypova, E., Zherebyateva, O., Tennikova, T., and Sarantseva, S. (2020) The neuroprotective effect of fullerenols on a model of Parkinson’s disease in Drosophila melanogaster. Biochem Biophys Res Commun. 523, 446–451

24. Golomidov, I. M., Latypova, E. M., Ryabova, E. V., Bolshakova, O. I., Komissarov, A. E., and Sarantseva, S. V. (2022) Reduction of the α-synuclein expression promotes slowing down early neuropathology development in the Drosophila model of Parkinson’s disease. J Neurogenet. 36, 1–10

25. Meulener, M. C., Xu, K., Thomson, L., Ischiropoulos, H., and Bonini, N. M. (2006) Mutational analysis of DJ-1 in Drosophila implicates functional inactivation by oxidative damage and aging. Proc Natl Acad Sci U S A. 103, 12517–12522

26. Miura, E., Hasegawa, T., Konno, M., Suzuki, M., Sugeno, N., Fujikake, N., Geisler, S., Tabuchi, M., Oshima, R., Kikuchi, A., Baba, T., Wada, K., Nagai, Y., Takeda, A., and Aoki, M. (2014) VPS35 dysfunction impairs lysosomal degradation of α-synuclein and exacerbates neurotoxicity in a Drosophila model of Parkinson’s disease. Neurobiol Dis. 71, 1–13

27. Suzuki, M., Fujikake, N., Takeuchi, T., Kohyama-Koganeya, A., Nakajima, K., Hirabayashi, Y., Wada, K., and Nagai, Y. (2015) Glucocerebrosidase deficiency accelerates the accumulation of proteinase K-resistant α-synuclein and aggravates neurodegeneration in a Drosophila model of Parkinson’s disease. Hum Mol Genet. 24, 6675–6686

28. Chen, L., and Feany, M. B. (2005) Alpha-synuclein phosphorylation controls neurotoxicity and inclusion formation in a Drosophila model of Parkinson disease. Nat Neurosci. 8, 657– 663

29. Nevzglyadova, O. V., Mikhailova, E. V., Artemov, A. V., Ozerova, Y. E., Ivanova, P. A., Golomidov, I. M., Bolshakova, O. I., Zenin, V. V., Kostyleva, E. I., Soidla, T. R., and Sarantseva, S. V. (2018) Yeast red pigment modifies cloned human α-synuclein pathogenesis in Parkinson disease models in Saccharomyces cerevisiae and Drosophila melanogaster. Neurochem Int. 120, 172–181

30. Ikeda, A., Nishioka, K., Meng, H., Takanashi, M., Hasegawa, I., Inoshita, T., Shiba-Fukushima, K., Li, Y., Yoshino, H., Mori, A., Okuzumi, A., Yamaguchi, A., Nonaka, R., Izawa, N., Ishikawa, K.-I., Saiki, H., Morita, M., Hasegawa, M., Hasegawa, K., Elahi, M., Funayama, M., Okano, H., Akamatsu, W., Imai, Y., and Hattori, N. (2019) Mutations in CHCHD2 cause α-synuclein aggregation. Hum Mol Genet. 28, 3895–3911

31. Periquet, M., Fulga, T., Myllykangas, L., Schlossmacher, M. G., and Feany, M. B. (2007) Aggregated alpha-synuclein mediates dopaminergic neurotoxicity in vivo. J Neurosci. 27, 3338–3346

32. Chen, L., Periquet, M., Wang, X., Negro, A., McLean, P. J., Hyman, B. T., and Feany, M. B. (2009) Tyrosine and serine phosphorylation of alpha-synuclein have opposing effects on neurotoxicity and soluble oligomer formation. J Clin Invest. 119, 3257–3265

33. Maor, G., Dubreuil, R. R., and Feany, M. B. (2023) α-Synuclein Promotes Neuronal Dysfunction and Death by Disrupting the Binding of Ankyrin to β-Spectrin. J Neurosci. 43, 1614–1626

34. Sarkar, S., Olsen, A. L., Sygnecka, K., Lohr, K. M., and Feany, M. B. (2021) α-synuclein impairs autophagosome maturation through abnormal actin stabilization. PLoS Genet. 17, e1009359

35. Shehadul Islam, M., Aryasomayajula, A., and Selvaganapathy, P. (2017) A Review on Macroscale and Microscale Cell Lysis Methods. Micromachines. 8, 83

36. Sawada, M., Yamaguchi, K., Hirano, M., Noji, M., So, M., Otzen, D., Kawata, Y., and Goto, Y. (2020) Amyloid Formation of α-Synuclein Based on the Solubility- and Supersaturation-Dependent Mechanism. Langmuir. 36, 4671–4681

37. Calvo, E., Bravo, R., Amigo, A., and Gracia-Fadrique, J. (2009) Dynamic surface tension, critical micelle concentration, and activity coefficients of aqueous solutions of nonyl phenol ethoxylates. Fluid Phase Equilibria. 282, 14–19

38. Matsuoka, K., and Moroi, Y. (2002) Micelle formation of sodium deoxycholate and sodium ursodeoxycholate (part 1). Biochim Biophys Acta. 1580, 189–199

39. Chern, C. S., Lin, S. Y., Chang, S. C., Lin, J. Y., and Lin, Y. F. (1998) Effect of initiator on styrene emulsion polymerisation stabilised by mixed SDS/NP-40 surfactants. Polymer. 39, 2281–2289

40. Pisárčik, M., Devínsky, F., and Pupák, M. (2015) Determination of micelle aggregation numbersof alkyltrimethylammonium bromide and sodiumdodecyl sulfate surfactants using time-resolvedfluorescence quenching. Open Chemistry. 13, 000010151520150103

41. Bandopadhyay, R. (2016) Sequential Extraction of Soluble and Insoluble Alpha-Synuclein from Parkinsonian Brains. J Vis Exp. 10.3791/53415

42. Iwatsubo, T., Yamaguchi, H., Fujimuro, M., Yokosawa, H., Ihara, Y., Trojanowski, J. Q., and Lee, V. M. (1996) Purification and characterization of Lewy bodies from the brains of patients with diffuse Lewy body disease. Am J Pathol. 148, 1517–1529

43. Campbell, B. C., Li, Q. X., Culvenor, J. G., Jäkälä, P., Cappai, R., Beyreuther, K., Masters, C. L., and McLean, C. A. (2000) Accumulation of insoluble alpha-synuclein in dementia with Lewy bodies. Neurobiol Dis. 7, 192–200

44. Campbell, B. C., McLean, C. A., Culvenor, J. G., Gai, W. P., Blumbergs, P. C., Jäkälä, P., Beyreuther, K., Masters, C. L., and Li, Q. X. (2001) The solubility of alpha-synuclein in multiple system atrophy differs from that of dementia with Lewy bodies and Parkinson’s disease. J Neurochem. 76, 87–96

45. Bandopadhyay, R. (2016) Sequential Extraction of Soluble and Insoluble Alpha-Synuclein from Parkinsonian Brains. J Vis Exp. 10.3791/53415

