## Supplementary figures and images for "Biochemical fractionation of human α-Synuclein in a *Drosophila* model of synucleinopathies"

### Supplemental Figure 1

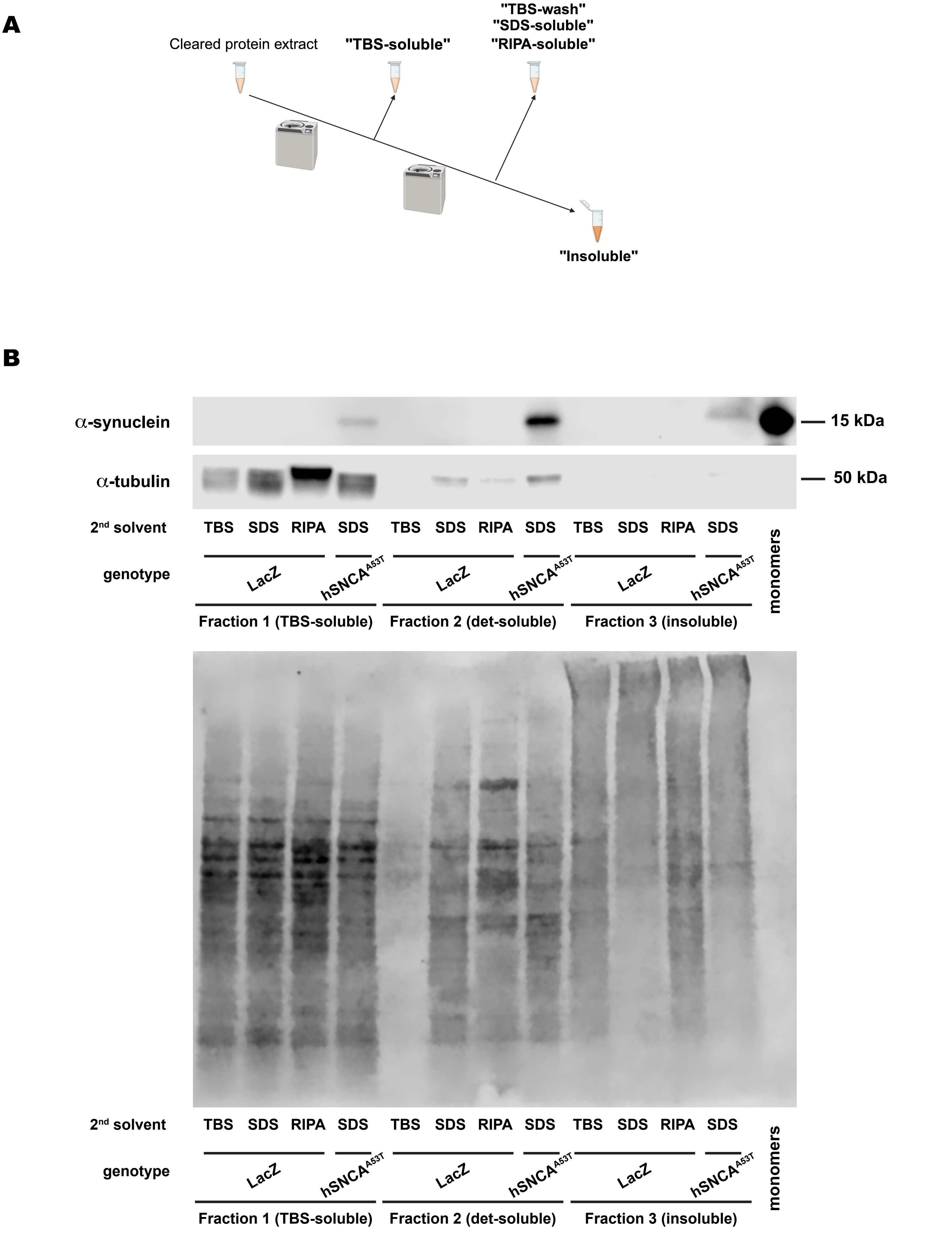

### Supplemental Figure 2

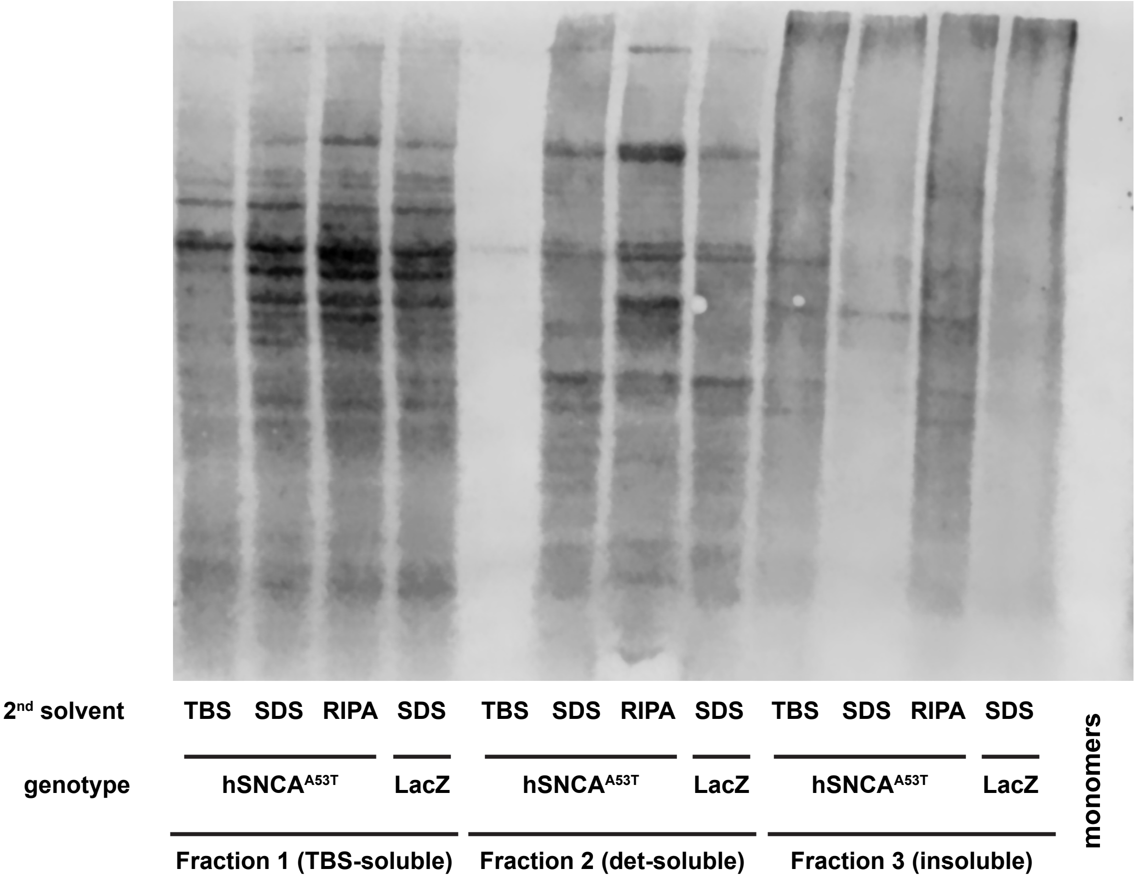

### Supplemental Figure 3

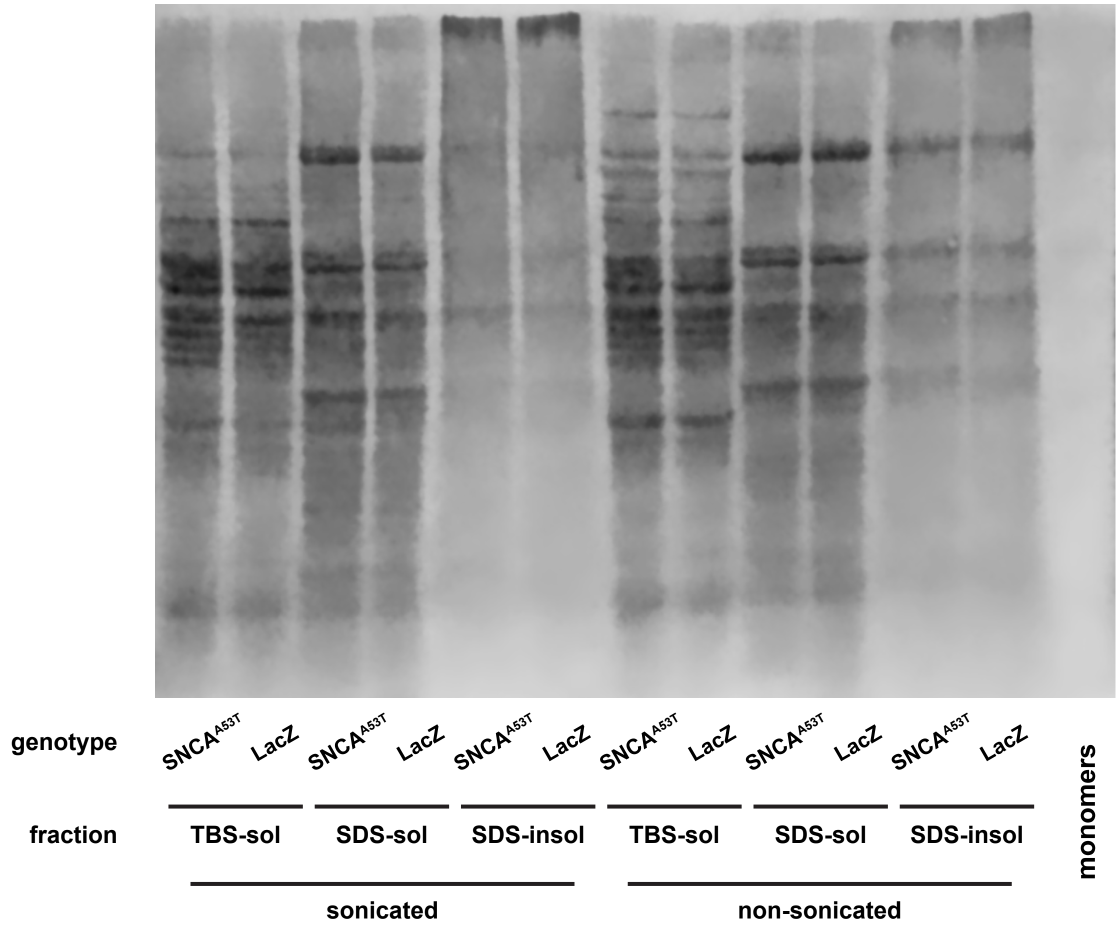

### Supplemental Figure 4

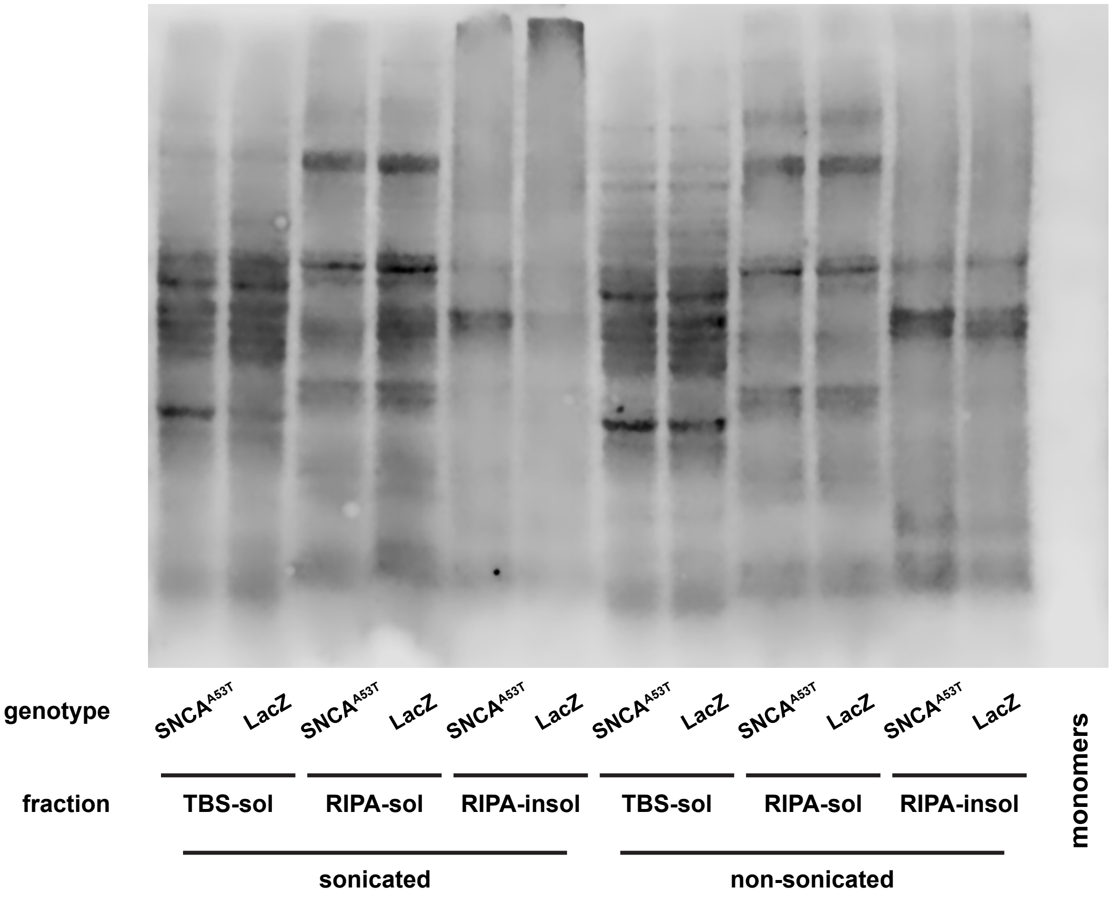

### Supplemental Figure 5

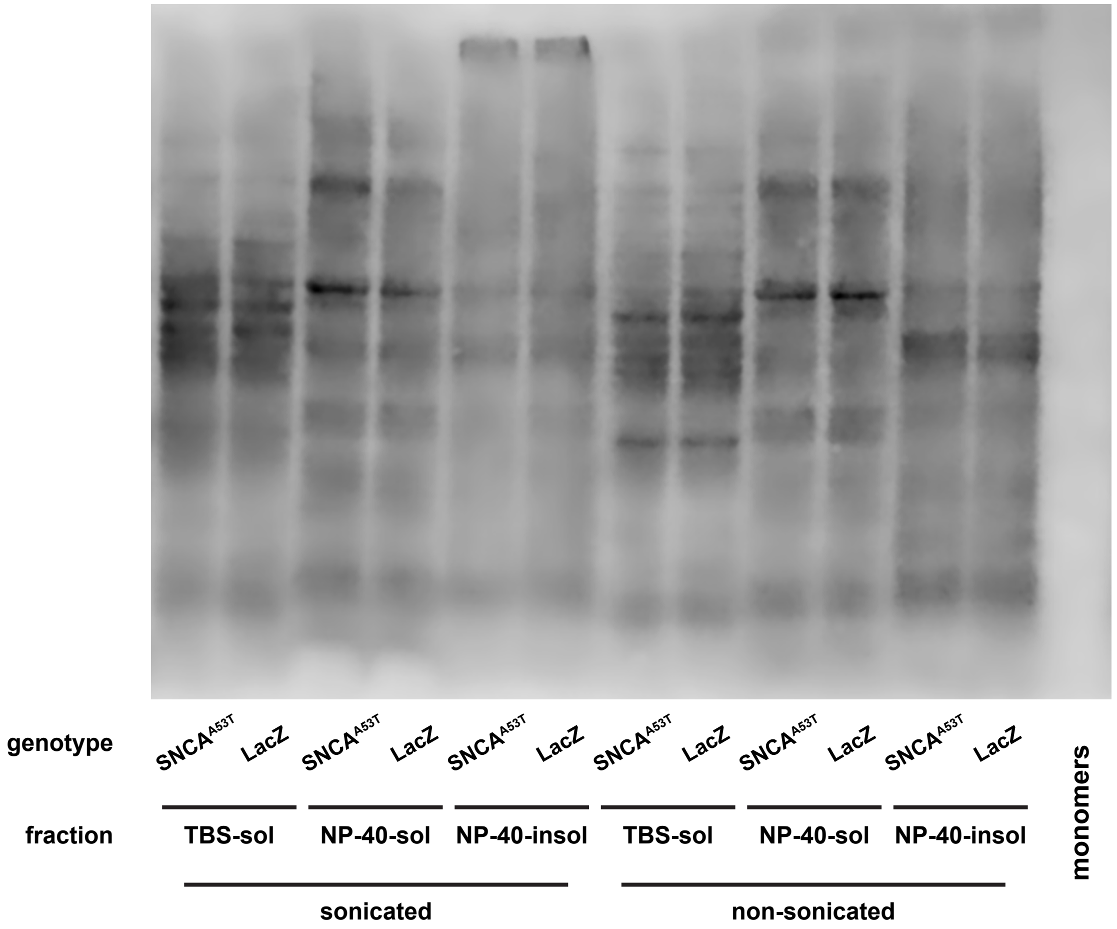

### Supplemental Figure 6

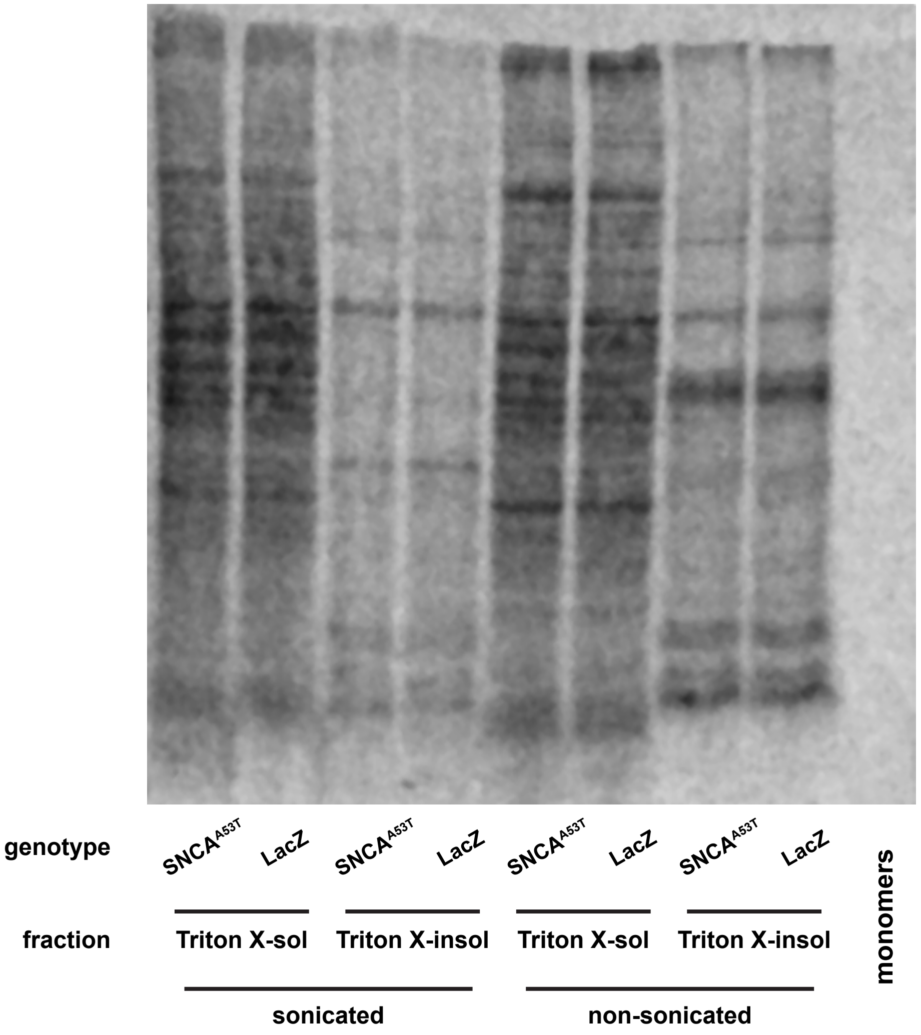
